# Determinants of Disordered Protein Co-Assembly Into Discrete Condensed Phases

**DOI:** 10.1101/2023.03.10.532134

**Authors:** Rachel M. Welles, Kandarp A. Sojitra, Mikael V. Garabedian, Boao Xia, Wentao Wang, Muyang Guan, Roshan M. Regy, Elizabeth R. Gallagher, Daniel A. Hammer, Jeetain Mittal, Matthew C. Good

**Author notes:** Equal contribution (Rachel M. Welles, Kandarp A. Sojitra).

## Abstract

Cells harbor numerous mesoscale membraneless compartments that house specific biochemical processes and perform distinct cellular functions. These protein and RNA-rich bodies are thought to form through multivalent interactions among proteins and nucleic acids resulting in demixing via liquid-liquid phase separation (LLPS). Proteins harboring intrinsically disordered regions (IDRs) predominate in membraneless organelles. However, it is not known whether IDR sequence alone can dictate the formation of distinct condensed phases. We identified a pair of IDRs capable of forming spatially distinct condensates when expressed in cells. When reconstituted in vitro, these model proteins do not co-partition, suggesting condensation specificity is encoded directly in the polypeptide sequences. Through computational modeling and mutagenesis, we identified the amino acids and chain properties governing homotypic and heterotypic interactions that direct selective condensation. These results form the basis of physicochemical principles that may direct subcellular organization of IDRs into specific condensates and reveal an IDR code that can guide construction of orthogonal membraneless compartments.

## Introduction

Cells compartmentalize biochemical processes to control reactivity and enforce selective interactions^1–3^. Inappropriate mixing of components, such as the presence of DNA in the cytosol or release of lysosomal proteases, damages the cell and triggers host defenses^4^. Membrane-bound compartments display restrictive permeability and enrichment, often requiring distinct targeting motifs^5, 6^. Macromolecules can also selectively partition into membraneless organelles, increasingly known as biomolecular condensates^7, 8^. These structures form via self-assembly and phase separation of proteins and RNAs into mesoscale compartments, demixed from the surrounding cellular milleu^9, 10^. Multivalent interactions along polypeptide chains with low complexity (LC) sequences or intrinsically disordered regions (IDRs) often drive demixing into condensed phases^11, 12^. Although dozens of cellular condensates have been characterized, it is known that a significant fraction of proteins in the human proteome have IDRs^13^ and thus it is not understood how thousands of intrinsically disordered proteins (IDPs) correctly partition into distinct subcellular condensates.

Selective assembly into orthogonal membraneless compartments could be directed by several mechanisms. It may be directed by IDR composition (shared residues among different IDRs), presence of folded oligomerization domains, or interactions with other macromolecules. Distinct condensates such as P-bodies and stress granules contain shared ligands, which can lead to condensate mixing^14^. The nucleolus is a multiphase structure containing immiscible protein phases^15^. NPM1 and FIB1 partition into distinct phases and their RNA binding domains - but not IDRs – likely explain their immiscibility within the nucleolus^16^. Additionally, RNA can alter co-condensation of FUS LC and Arg-rich peptides, resulting in distinct phases^17^. However, a major gap in our knowledge is whether selective partitioning is specified directly by primary sequence and/or the chain properties of disordered polypeptides. To ask simply, to what extent is it possible for two IDRs to selectively assemble into distinct condensates with minimal co-mixing in the absence of shared ligands or macromolecules?

Recently, our lab and others have revealed an amino acid code governing homotypic interactions that underlie liquid-liquid phase separation (LLPS) of various IDRs^18–23^. Coarse-grained models have provided insights on key residues or motifs that influence IDR saturation concentrations (C_sat_) and client partitioning to condensed phases. However, it has not been possible to predict whether heterogenous mixtures of IDPs will co-condense or form distinct condensates^24^. A fundamental challenge is that IDRs lack secondary structure and are often amphipathic, with a near-full repertoire of amino acid chemistries and shared molecular interactions via common backbone atoms. For example, a subset of prion-like domains (PLDs) co-partition in transcriptional condensates^25^. Analysis of protein blends having multivalent interactions, while instructive, cannot explain whether or how selective condensation of IDRs is achieved^26^. Further, biochemical reconstitution may not fully recapitulate selectivity as distinct condensed phases ex vivo may in fact co-partition in the complex cytosol of a cell.

We sought to determine the extent to which IDPs selectively partition into discrete condensed phases and to determine the chemical principles of selectivity (Fig. 1A). To simplify identification of polypeptide sequences that are orthogonal to one another with respect to LLPS, we chose to express pairs of IDRs and full-length (FL) proteins in a cellular context that minimizes interactions with endogenous proteins and more directly assesses selectivity of condensates alone. We tested inducible expression of these proteins fused to fluorophores and assessed demixing into spatially distinct versus mixed condensed phases in cytosol. After identifying a range of LLPS specificity, we focused our study on a pair of IDRs: FUS LC^27^ and LAF-1 RGG^28^. We found that this pair forms discrete condensates that do not co-partition when reconstituted in vitro. Thus, condensate specificity in this case is hard-coded in protein sequence and does not require additional factors such as RNA or ligands. By combining experiment with molecular simulation, we identified chain-level properties controllable via changes in amino acid sequence that govern the specificity of their condensation into distinct phases. Specifically, our data are consistent with a mechanistic model in which competition between homotypic and heterotypic interactions of an IDR ultimately dictates partitioning. We propose that these rules can inform how IDPs may partition in cells and can be generalized to additional sets of IDPs enabling formation of distinct supramolecular assemblies in cells for applications in synthetic biology^29^.

**Fig. 1.**
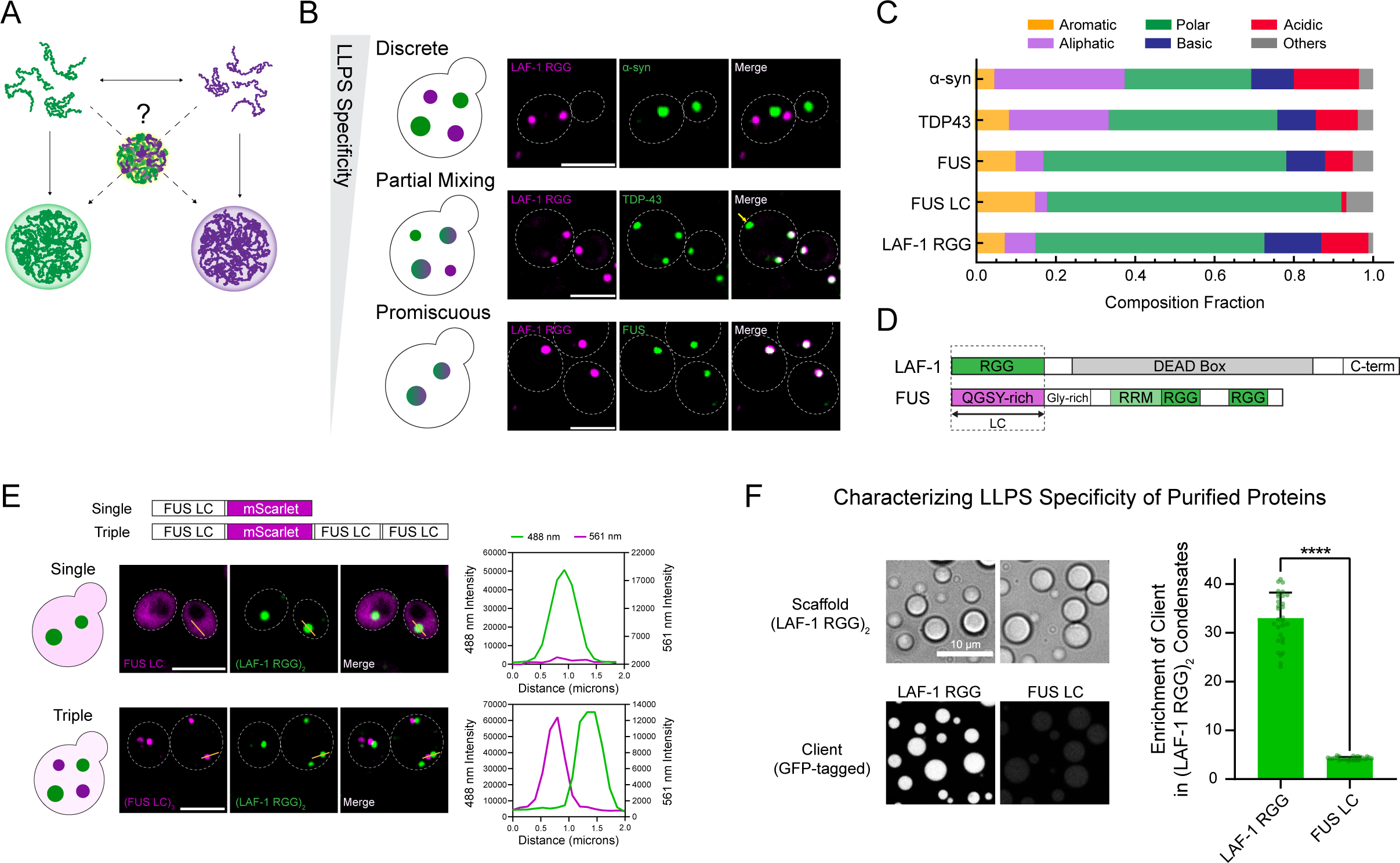
Identification of disordered proteins that form discrete condensed phases in living cells and when biochemically reconstituted in vitro. A. Schematic of central question: are two disordered proteins capable of forming distinct condensates or will they co-partition upon demixing? B. Co-expression of pairs of IDPs in model eukaryotic cell, *S. cerevisiae*, reveals a range of LLPS specificity. LAF-1 RGG expressed as tandem fused to mScarlet. Co-expressed IDPs α-synuclein, TDP-43, and FUS are fused to GFP. Cells imaged after overnight induction of IDP expression. Scale bars: 5 μm. C. Plot showing amino acid composition of IDPs tested in 1B. D. LAF-1 and FUS contain shared domains, RGG, RRM that likely promote co-partitioning. However, each also has an approximately 160 amino acid long IDR or low complexity (LC) domain. E. FUS LC partitioning in cells containing condensates formed by expression of LAF-1 RGG. Scale bars: 5 μm. FUS LC-mScarlet does not condense or partition into (LAF-1 RGG)_2_-GFP condensates. A version with three repeats of IDR (FUS LC)_3_-mScarlet is capable of forming condensates and does not co-partition with (LAF-1 RGG)_2_-GFP condensates. F. Purified forms of LAF-1 RGG and FUS LC show selective partitioning to preformed condensates of (LAF-1 RGG)_2_. Quantitation of enrichment of GFP-tagged forms of LAF-1 RGG and FUS LC in condensates, compared to continuous phases. n = 30 condensates per column. Scale bar: 10 μm. Error bars represent 95% confidence intervals. Significance was calculated by a Welch’s t-test; ns P > 0.05, * P ≤ 0.05, ** P ≤ 0.01, *** P ≤ 0.001, and **** P ≤ 0.0001.

## Results

### Characterization of IDP orthogonality and formation of discrete condensed phases

To determine the extent to which IDPs are capable of selective partitioning into spatially distinct condensed phases, we co-expressed pairs of IDPs with or without folded domains and fused to different fluorescent tags in the model eukaryote, budding yeast. We reasoned that expression of metazoan IDPs in a simplified eukaryotic cytosol, largely absent from their endogenous interaction networks, could potentially reveal LLPS specificity attributable to the individual IDP sequences. Further, *Saccharomyces cerevisiae* has been used extensively as a model cell to study protein aggregation and protein condensation, revealing rules of their genetic and chemical regulation^30–36^.

We compared co-condensation versus partitioning of a handful of well-characterized IDRs or full-length proteins after inducible expression and identified a range of LLPS selectivity (Fig. 1B). We categorized protein pairs that fell into three scenarios: discrete compartments, partial mixing, and promiscuous co-partitioning. Co-expression of α-synuclein FL and LAF-1 RGG, led to fully orthogonal foci. Co-expression of TDP-43 FL and LAF-1 RGG led to overlapping condensates and some discrete phases. FUS FL and LAF-1 RGG almost completely colocalize (Fig. 1B, quantitation in Extended Data Fig. S1A). Despite this range of specificity, the amino acid compositions of these proteins are actually quite similar (Fig. 1C). However, because we used full length TDP-43 and FUS proteins, it was possible that promiscuity could be attributed to additional domains outside the IDRs. For example, FUS contains RGG and RRM domains that could interact with LAF-1 RGG (Fig. 1D). Therefore, we tested co-expression of the low-complexity domain of FUS (FUS LC) with LAF-1 RGG. The FUS LC domain has a C_sat_ of ∼100 μM^22, 37^ and, thus, does not form condensates when expressed in cells, allowing us to measure its co-partitioning as a client into the LAF-1 RGG condensed phase. Interestingly, FUS LC does not enrich within LAF-1 RGG condensates (Fig. 1E). To ensure that this specificity was not related to its inability to condense, we also expressed a FUS LC trimer capable of condensation due to a reduced C_sat_^18^ and found it forms spatially distinct condensates from LAF-1 RGG (Fig. 1E, quantitation of EI in Extended Data Fig. S1B). These data indicate that LLPS specificity of this IDR pair may be related to the differences in their protein sequence composition. Coarse-grained simulations of an equimolar binary solution of LAF-1 RGG and FUS LC further indicated formation of distinct phases (Extended Data Fig. S1C). To determine whether LLPS specificity was indeed encoded in protein sequence, we purified these IDRs and reconstituted their assembly into liquid-like condensates in vitro. We formed droplets of LAF-1 RGG and measured the recruitment of fluorescently labeled ‘clients’ at concentrations well below their C_sat_, such that the clients could not condense on their own. We observed strong partitioning of GFP-tagged LAF-1 RGG but not GFP-tagged FUS LC within LAF-1 RGG condensates (Fig. 1F), suggesting that these IDR sequences directly encode chemical determinants for selective condensation. Taken together, these results suggest that native IDRs can form discrete condensed phases and this orthogonal IDP pair afforded us a platform to uncover residues governing specificity.

### Amino Acid Sequence Determinants of FUS LC and LAF-1 RGG Condensation Specificity

#### Contribution of Charged Residues to Selective LLPS

Polypeptide sequence features governing homotypic phase separation of a handful of individual IDPs have recently been characterized^12, 20, 22, 38–40^. Residues that determine the C_sat_ of FUS have been described^41, 42^ and, separately, the amino acids and chain properties that dictate the C_sat_ of LAF-1 RGG have been identified^19, 20^. The sequences of FUS LC and LAF-1 RGG domains differ in composition (Fig. 2A), which provided us a means to interrogate the specific residues that control the LLPS specificity of FUS LC^43^. FUS LC largely lacks charged residues: it has no basic and only two acidic residues (Fig. 2A). In contrast, LAF-1 RGG contains 24 basic residues (all Arg), and 20 acidic residues. We first tested whether addition of charged residues to FUS LC would promote co-partitioning with LAF-1 RGG condensates. We quantify the amount of protein in a condensate through a partitioning coefficient or enrichment index (EI), the ratio of intensity in the condensate to intensity in the cytoplasm. As noted above, because FUS LC has a high C_sat_, it cannot condense on its own but its EI can be measured as a client within LAF-1 RGG condensates. Our experimental setup for co-expression in yeast throughout the paper (unless noted otherwise) uses tandem condensing LAF-1 RGG, (LAF-1 RGG)_2_-GFP, as a scaffold, and non-condensing single FUS LC constructs attached to mScarlet as clients. In terms of range of possible enrichment in LAF-1 RGG tandem condensates, we note that the maximum enrichment possible for a single IDR as a client is achieved via LAF-1 RGG-mScarlet, with an EI of 6.57. In contrast, no enrichment is an EI of 1, which is similar to the EI of 1.19 we measure for FUS LC-mScarlet in LAF-1 RGG condensates (Extended Data Sig. S1B). We assayed the effect on specificity of adding charged resides to FUS LC in place of serine residues. Surprisingly, neither addition of 16 balanced charges (‘Charge+16’: 16 Ser → 8 Arg + 8 Asp) nor addition of 32 balanced charges (‘Charge+32’: 32 Ser → 16 Arg + 16 Asp) had any effect on FUS LC partitioning within LAF-1 RGG condensates (Fig. 2B, C). FUS LC mutants were well-mixed in the cytoplasm and maintained their selectivity, with no significant enrichment within LAF-1 RGG foci (Extended Data Fig. S2A).

**Fig. 2.**
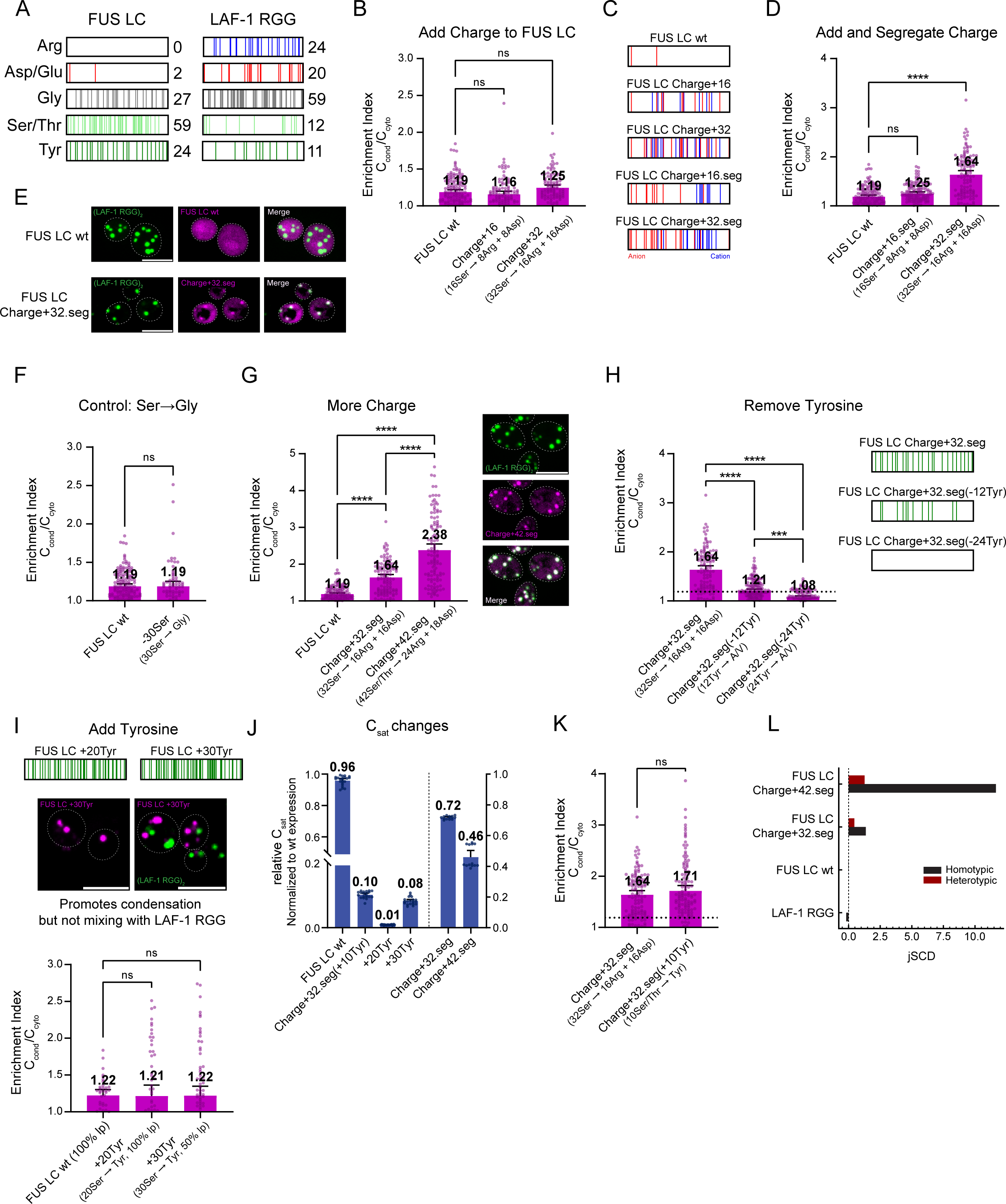
Amino acid sequence determinants of FUS LC coacervation specificity. A. Positions and counts of amino acid subtypes in the IDRs from FUS LC wildtype and LAF-1 RGG. (B-K) Co-expression of FUS LC or mutant forms of it fused to mScarlet in cells containing condensates of (LAF-1 RGG)_2_-GFP. B. Addition of 16 or 32 charges to FUS LC, replacing Ser with equal portions of Arg and Asp, is not sufficient to cause FUS LC to partition to LAF-1 RGG condensates. N ≥ 108 condensates per column. C. Positions of charged residues in FUS LC wildtype and 4 charged variants. D, E. Addition and segregation of 32 charges (‘Charge+32.seg’) causes FUS LC to begin to partition to LAF-1 RGG condensates; it becomes promiscuous. Quantitation of FUS LC enrichment from confocal images of cells co-expression (E). n ≥ 115 condensates per column. F. Mutation of 30 Ser→Gly on FUS LC does not promote partitioning to LAF-1 RGG condensed phase. n ≥ 74 condensates per column. G. Addition and segregation of 42 charges promotes very strong co-mixing of FUS LC into LAF-1 RGG condensates. Confocal images (right). Overlap ‘merge’ is white. n ≥ 115 condensates per column. H. Removal of Tyr residues from FUS LC Charge+32.seg reduces its partitioning into LAF-1 RGG condensates. Location of Tyr residues in mutants shown to right of plot. N ≥ 124 condensates per column. I. Addition of Tyr to FUS LC is sufficient to promote its own condensation but not partitioning to LAF-1 RGG condensates. Top row shows location of Tyr residues in mutants. n ≥ 37 condensates per column. J. Estimation of relative C_sat_ for each FUS LC mutant from cells with visible condensates, normalized to expression level of FUS LC wildtype on day of imaging. n between 13 and 22 for all columns. K. Addition of 10 Tyr residues to Charge+32.seg does not significantly increase mixing with LAF-1 RGG condensates. n ≥ 115 condensates per column. L. Homotypic and heterotypic jSCD^81^ values of FUS LC wildtype and charged segregated mutants. jSCD parameter is useful to predict binding affinity of two components and can be used to estimate changes in the homotypic and heterotypic interactions between two IDPs upon changes in charge patterning. Error bars represent 95% confidence intervals. Significance was calculated by one-way analysis of variance (ANOVA) except for F and J which utilized a Welch’s t-test; ns P > 0.05, * P ≤ 0.05, ** P ≤ 0.01, *** P ≤ 0.001, and **** P ≤ 0.0001. Scale bars: 5 μm.

Patchy or segregated charge on IDPs can promote homotypic self-assembly, lowering the C_sat_ for condensation^20, 44, 45^. Thus, we tested whether segregation of these additional charged residues on FUS LC, which were relatively well-mixed in the ‘Charge+16’ and ‘Charge+32’ mutants, would promote its partitioning with LAF-1 RGG. Only the FUS LC mutant ‘Charge+32.seg’ (32 Ser → 16 Arg + 16 Asp, segregated; Fig. 2C) significantly enriched within LAF-1 RGG condensates (Fig. 2D, E). To check whether this was related to the loss of serines, we tested FUS LC mutant ‘-30Ser’ (30 Ser → Gly) (Fig. 2F). In co-expression, it showed no change from FUS LC wildtype with the LAF-1 RGG condensates, suggesting that gain and segregation of charged residues specifically led to the promiscuous interactions between FUS LC and LAF-1 RGG in case of ‘Charge+32.seg’ mutant. In support of this, addition of more segregated charge in FUS LC (‘Charge+42.seg’: 42Ser/Thr → 24 Arg, 18 Asp, segregated) led to even stronger partitioning into LAF-1 RGG condensates (Fig. 2G). This charge addition matches the exact number of charged residues on LAF-1 RGG, enriching 2-3 times higher than that of FUS LC wildtype. Notably, the co-expression of FUS LC and its mutants were similar and had little effect on the condensation of LAF-1 RGG. The average number of LAF-1 RGG condensates across all co-expressed FUS LC mutants remained within twofold of FUS LC wildtype, indicating no significant disruption to scaffold condensation (Extended Data Fig. S2B). The levels of FUS LC were only modestly impacted by mutation; each variant reached an intracellular concentration within approximately two-fold of wildtype (Extended Data Fig. S2C).

To determine the mechanism behind the loss of specificity due to charge segregation, we purified FUS LC mutants Charge+32.seg and Charge+42.seg and formed condensates in vitro (Extended Data Fig. S2D-G). FUS LC wildtype condensates showed minimal enrichment of fluorescent (LAF-1 RGG)_2_-GFP tracer (Extended Data Fig. S2D, E). In contrast, condensates of FUS LC mutant Charge+32.seg strongly enriched (LAF-1 RGG)_2_-GFP. Condensates of FUS LC mutant Charge+42.seg also enriched (LAF-1 RGG)_2_-GFP to a larger extent than condensates formed by FUS LC wildtype, though its EI is less than that of Charge+32.seg. Sequence charge decoration (SCD)^46^ is a parameter that quantifies the degree of charge patterning within a sequence. Sequences with highly segregated opposing charges will have more negative SCD scores and a lower C_sat_ for phase separation^20^. This is the case for the surfactant activity of protein Ki-67^47^ and for raising the transition temperature (T_t_) for LLPS of LAF-1 RGG sequence shuffles^20^. Indeed, FUS LC mutants Charge+32.seg and Charge+42.seg both form condensates at about two orders of magnitude lower concentrations than FUS LC wildtype (Extended Data Fig. S2F). Additionally, turbidity assays reveal that purified FUS LC mutant Charge+32.seg has a greatly elevated T_t_ compared to wildtype, and that FUS LC mutant Charge+42.seg has an even higher T_t_ (Extended Data Fig. S2G). Overall, these data suggest that the near absence of charged residues contributes to the LLPS specificity of the FUS LC IDR. However, addition of charges to FUS LC alone cannot promote mixing with LAF-1 RGG; these charges must be segregated or blocky and the effect depends non-monotonically on the number of charged residues added.

### Contributions of Tyrosine to LLPS Selectivity

Homotypic condensation of LAF-1 RGG also depends on both Arg and Tyr residues along the polypeptide chain^20, 39^. Therefore, we wondered what role Tyr in FUS LC plays in selective LLPS or, conversely, heterotypic partitioning with LAF-1 RGG. We interrogated the role of Tyr in two ways: What is the effect of removal of Tyr from FUS LC mutant, Charge+32.seg, which promiscuously partitions to the LAF-1 RGG condensed phase? Does addition of more Tyr to FUS LC wildtype, which does not mix with LAF-1 RGG, lead to loss of specificity? We started by removing half or all 24 Tyr from Charge+32.seg. When Charge+32.seg(-12Tyr) and Charge+32.seg(-24Tyr) were co-expressed with LAF-1 RGG, we observed a stepwise drop in their enrichment in LAF-1 RGG condensates (Fig. 2H). Charge+32.seg(-24Tyr) has lower enrichment values even than wildtype. These results demonstrate that Tyr residues contribute to heterotypic interactions between FUS LC and LAF-1 RGG. To assess whether Tyr residues alone are adequate to increase coacervation with LAF-1 RGG, we created FUS LC mutants with 20 and 30 additional tyrosines (‘+20Tyr’ and ‘+30Tyr’). When expressed alone or co-expressed with LAF-1 RGG, these FUS LC mutants with added Tyr were now capable of condensing on their own in yeast cells (Fig. 2I), something not seen for prior mutants. This is consistent with previous studies correlating increased Tyr content to lowered IDR C_sat_^21^. Expression of tyrosine-rich mutants was dramatically reduced in yeast (Extended Data Fig. S2C). Despite this, their C_sat_ in cells was far lower than FUS LC wildtype, thereby promoting their condensation (Fig. 2J). Surprisingly, neither mutant (+20Tyr or +30Tyr) showed significant enrichment with LAF-1 RGG (Fig. 2I), which may have been expected based on the Tyr removal data (Fig. 2H), suggesting that Tyr alone in this context cannot dictate co-partitioning. Furthermore, addition of 10 Tyr to Charge+32.seg (Charge+32.seg(+10Tyr)) also lowered C_sat_ but had no significant effect on its co-partitioning with LAF-1 RGG condensates (Fig. 2K). Consistent with in vitro data, the relative in vivo C_sat_ for the FUS LC mutants containing added charge, Charge+32.seg and Charge+42.seg, decreased relative to FUS LC wildtype, but somewhat surprisingly were approximately an order of magnitude higher than the mutants with added Tyr (Fig. 2J). These results hint at the potential role of homotypic interactions in addition to the obvious contribution of heterotypic interactions in controlling co-partitioning of IDRs. It is natural to expect that residues predicted to promote interactions between different proteins can also strengthen homotypic interactions, thereby either enhancing or blocking co-partitioning, which is dependent upon the competition between homotypic and heterotypic interactions. For example, addition of 10 charges to Charge+32.seg increases predicted heterotypic interactions, as measured by joint SCD (jSCD)^48^, but also promotes a larger increase in homotypic interactions (Fig. 2L). Due to challenges of independently controlling homotypic and heterotypic interactions experimentally, we utilized a minimal computational model to better understand our findings.

### Minimalistic polymer model for two-component phase separation

Experimental results so far clearly highlight the interplay of chain-level homotypic and heterotypic interactions instead of just the presence or absence of certain amino acids in controlling the specificity of discrete condensate formation and in the co-partitioning of two IDRs. Previous studies on protein LLPS, including complex multicomponent phase behavior of mixtures, have successfully used simple computational models guided by the essential polymer physics principles^49–54^. Many of these studies highlighted the role of specific heterotypic interactions between multiple components that can be controlled independently from homotypic interactions in many systems such as polySUMO-polySIM, polySH3-polyPRM, PTB-RNA, and SynGAP-PSD95 in the stability of condensates formed via scaffold-client interactions, client valency, and the scaffold-client stoichiometry^17, 26, 55–60^. However, results based on these earlier studies cannot explain the scenarios that we observed in co-partitioning of FUS LC mutants and LAF-1 RGG. For example, we do not observe enhanced enrichment with the addition of 20 and 30 Tyr to FUS LC (‘+20Tyr’ and ‘+30Tyr’), which should enhance heterotypic interaction between the two but also change the homotypic interactions between FUS LC mutants significantly. To better understand our experimental findings, we used a simple homopolymer model for representing LAF-1 RGG and FUS LC as described next (see Methods for more details).

We used two types of polymers, A and B, where A acts as a scaffold (LAF-1 RGG) and B acts as client (FUS LC). Interactions between monomers of each polymer with itself (λ_AA_ and λ_BB_) and monomers of other type (λ_AB_) can be tuned independently in our model and are normalized with respect to the homotypic interactions between polymer A’s monomers, i.e., λ_AA_ = 1. Due to the high LLPS propensity of the LAF-1 RGG scaffold in our experiment, we varied the interactions λ_BB_ and λ_AB_ from 0 to 1 (Fig. 3A). We performed standard coexistence simulations in the slab geometry of an equimolar mixture of A and B polymers in which A polymer forms a liquid-like condensed phase and the B polymer concentration changes inside A’s condensed phase with increasing λ_BB_ and/or λ_AB_ ^61^. Similar to the analysis of experimental data, we computed enrichment index (EI) as the ratio of B’s concentration inside A to its solution concentration (and not dilute phase concentration due to numerical issues at low values) to quantify the partitioning of B polymer inside A polymer condensate. We note that for the rest of the discussion in this paper, we focus on input control parameters λ_BB_ and λ_AB_ for interpreting our results instead of explicit considerations of energetic and entropic contributions separately. It is worth pointing out that recent work on homotypic and heterotypic condensation in systems with specific one-to-one interactions (unlike promiscuous interactions between IDRs) have highlighted the role of entropic changes as the driving force for condensation^53, 57^. Our MD simulations of the minimalistic model explicitly account for any changes in entropy associated with the polymer chains between the dilute and condensed states (see Supplementary Figure S1 and text). We focus on the equilibrium behavior governed by minimum in free energy while taking the liberty to explain the observed EI changes in terms of homotypic and heterotypic interactions between the monomers, i.e., λ_BB_ and λ_AB_, respectively. This is a reasonable approach considering the existence of entropy-enthalpy compensation in many biomolecular assembly processes^62^, though future work should directly test this by conducting a detailed thermodynamic analysis.

**Fig. 3.**
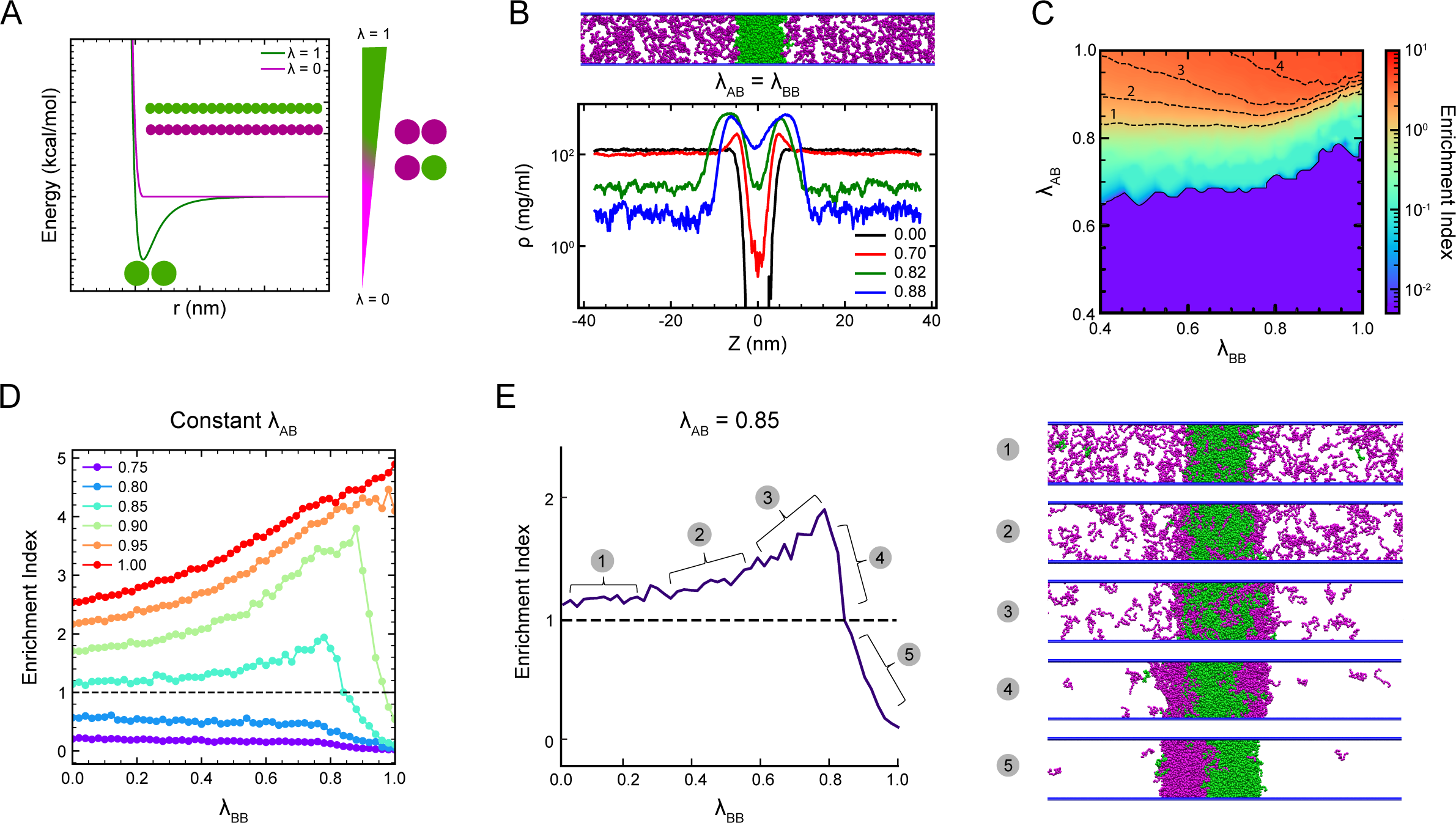
Minimalistic polymer model uncovers co-phase separation rules. A. Description of the homopolymer models and associated nonbonded pair potential (Methods). Interactions between monomers of polymer A (green) are fixed (λ = 1), while interactions between monomers of polymer B (magenta) and the heterotypic interaction between A and B monomers varied from λ = 0 to 1. B. Simulation snapshot of 250 chains each of A (green) and B (magenta) and concentration profiles of polymer B with respect to polymer A condensed phase centered in the middle of the simulation box. Black and red line corresponds to no (EI ∼ 0) or very low co-partitioning (EI << 1), while green line corresponds to onset of co-partitioning (EI ∼ 1), and blue line shows enhanced co-partitioning (EI > 1). C. Colormap of EI from coexistence scanning simulation with varying λ_AB_ (y-axis) and λ_BB_ (x-axis). Black dashed line corresponds to the contour line at constant EI values of 1, 2, 3, and 4. D. Plot showing variation of EI as a function of λ_BB_ at different constant λ_AB_ values (see legend). For high heterotypic interaction (λ_AB_ = 0.85 and 0.90), increasing B’s homotypic interaction enhances enrichment and an optimum homotypic interaction is required for maximum co-partitioning. When B’s homotypic is stronger than heterotypic interaction, enrichment decreases. E. Cartoon representation of two component system corresponding to various regimes in EI curve at λ_AB_ = 0.85.

Interestingly, the concentration of B polymers is very low inside A’s condensate until λ_BB_ = λ_AB_ = 0.82 (Extended Data Fig. S3A), where it reaches a similar value as the dilute phase and the solution concentration, i.e., EI ∼ 1 (Fig. 3B). This would suggest that FUS LC wildtype and LAF-1 RGG interactions are weakly attractive under experimental conditions, as EI close to 1 is observed, unlike partitioning of synthetic polymers such as PEG in aqueous two-phase systems in which EI << 1 ^63, 64^. Elevated concentrations of B at the interface for non-zero λ_AB_ reflect locally enhanced concentration of client molecules due to favorable heterotypic interactions, which are not sufficiently strong to cause co-mixing but may be sufficient to alter condensates’ material properties and composition control via a thin interfacial layer^65–68^.

Extensive simulations as a function of λ_BB_ and λ_AB_ were conducted (see Methods) to obtain EI over the whole range of homotypic and heterotypic interactions (Fig. 3C, Extended Data Fig. S3B). As expected, low values of homotypic and heterotypic interactions yield EI < 1 with a nonlinear change in EI with increasing λ_BB_ and λ_AB_. Even though strong heterotypic attractions alone (λ_BB_ = 0) are sufficient to cause co-condensation via complex coacervation (Extended Data Fig. S3C), changing homotypic interactions can cause a significant change in EI even for fixed λ_AB_ (Fig. 3D). Most importantly, a non-monotonic change in EI is observed with increasing λ_BB_ at fixed λ_AB_, which suggests that strengthening homotypic interactions between client polymers can initially enhance co-mixing, but there is an eventual rapid turnover to lower EI values upon a significant increase of λ_BB_. Interestingly, stronger heterotypic interactions (λ_AB_) can shift λ_BB_ to higher values at which EI turnover is observed, reflecting the complex interplay between the two. When we plot the ratio of heterotypic and homotypic interactions at the maximum EI, the ratio is approximately one (Extended Data Fig. S3D), which implies that for enrichment, homotypic interactions between client proteins should be similar or weaker than scaffold-client heterotypic interactions.

To understand and relate these general observations from our computational model to the experimental results in the previous sections, we look at the simulation snapshots and associated concentration profiles of A and B polymers (Fig. 3E and Extended Data Fig. S3E). Initially, at a high λ_AB,_ increasing λ_BB_ will enhance interactions among B polymers which leads to its increased recruitment inside A’s condensate. Interestingly, we find that large clusters of B polymers start to appear in the simulation box when approaching the maximum EI value, suggesting its homotypic condensation. Beyond λ_BB_ ∼ 0.8, B polymer forms a stable condensed phase resulting in a significant drop in co-partitioning or EI. To further probe the connection between B polymer’s LLPS propensity and its co-condensation with A polymer, we also checked the single-component LLPS behavior of B polymers with increasing λ_BB_. We find that polymer B alone forms a stable condensed phase for λ_BB_ ≥ 0.75 (Extended Data Fig. S3F).

Next, we use the above results from the minimal model to explain the observed experimental scenarios (Fig. 3E). FUS LC wildtype does not co-partition with LAF-1 RGG (Fig 3E, cartoon 1). As we add charges and segregate them (Charge+32.seg), it will enhance both homotypic and heterotypic interactions and thus shows enhanced co-partitioning (cartoon 2). With the further addition of charges, EI will increase further (cartoon 3). For FUS LC mutants where more Tyr are added (‘+20Tyr’ or ‘+30Tyr’), the homotypic interactions are strong enough to cause homotypic condensation under experimental conditions due to its very low C_sat_ (Fig. 2J), resulting in reduced co-partitioning within the LAF-1 RGG condensate. Instead, the experimental results exhibit similar EI values before and after the addition of Tyr in FUS LC wildtype or Charge+32.seg mutant. Two potential explanations for this apparent discrepancy can be considered. (1) One possibility is that the change in homotypic and heterotypic interactions upon these mutations (which are not feasible to quantify experimentally) can yield similar EI values on both sides of the maximum (Extended Data Fig. S3G). (2) It is also possible that the simulations conducted with a fixed number of finite chains (canonical ensemble) instead of a constant chemical potential overestimate the depletion of polymer B chains from A’s condensate upon homotypic condensation of B, which is reflected in the sharp drop in EI curves after reaching the maximum value. It will be worth testing this using on-lattice polymer models in future, which can be used to conduct simulations in a grand canonical ensemble (constant chemical potential)^69^.

In addition to providing a framework to understand the observed co-partitioning of LAF-1 RGG and FUS LC mutants, our computational model results should further be helpful in tuning FUS LC partitioning by changing the extent of homotypic and heterotypic interactions, for example, by leveraging the multivalency of repeat proteins.

### Polypeptide Chain Properties Governing FUS LC Partitioning and LLPS

#### IDR length

We next sought to determine - separate from amino acid sequence - whether alterations to polypeptide chains of FUS LC or LAF-1 RGG would experimentally impact their ability to form distinct condensed phases. We first tested whether lengthening the FUS LC IDR sequence would be sufficient to promote co-partitioning with LAF-1 RGG condensates (Fig. 4A). However, tandem FUS LC showed no enhanced interaction as measured by enrichment with co-expressed LAF-1 RGG (Fig. 4B). In contrast, for the Charge+32.seg FUS LC mutant, we observed that doubling of chain length significantly increased co-partitioning to the LAF-1 RGG condensed phase (Fig. 4B, C). These results demonstrate that increasing IDR length can increase the strength of existing multivalent interactions, but is not sufficient to broadly promote co-mixing, which is also highlighted by the results of the computational model in the limit of weak homotypic and heterotypic interactions. Specifically, simulation results with varying polymer lengths show relatively little change in co-partitioning for low homotypic and heterotypic attraction (Extended Data Fig. S4). But if heterotypic and/or homotypic interactions are stronger (e.g., Charge+32.seg FUS LC mutant), significant enhancement in EI can be observed, as long as homotypic interactions between client polymers are not strong enough to cause condensation (Fig. 4D).

**Fig. 4.**
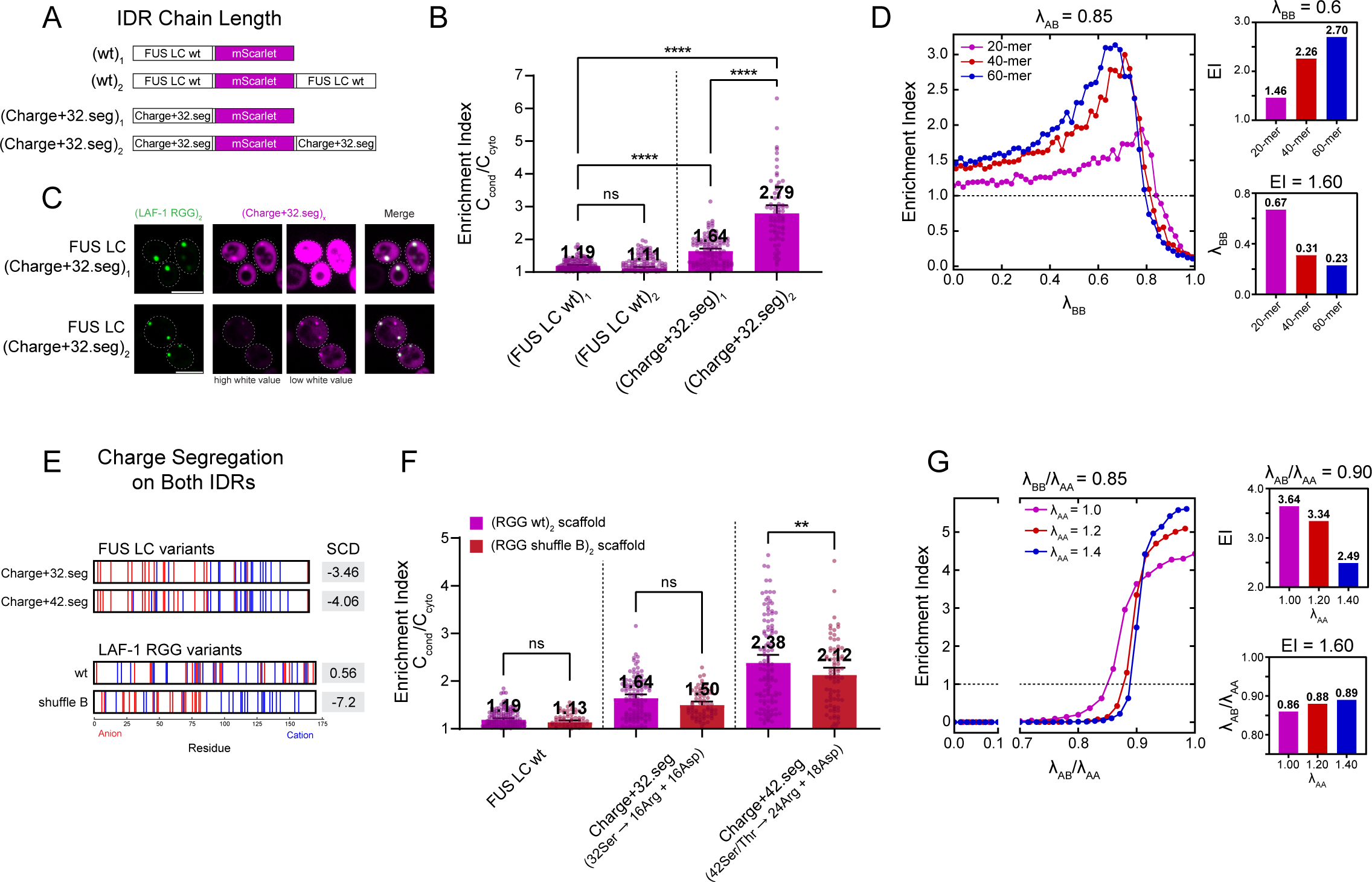
Polypeptide chain properties governing FUS LC partitioning and LLPS. A. Schematic: varying number of FUS LC wildtype or mutant IDRs fused to mScarlet. B. Quantitation of 2B. Tandem FUS LC wildtype does not partition to LAF-1 RGG condensates. Tandem FUS LC Charge+32.seg partitions even more strongly to LAF-1 RGG condensates than Charge+32.seg single, indicating a role for IDR chain length in LLPS specificity. n ≥ 68 condensates per column. C. Confocal images of cells expressing FUS LC mutant Charge+32.seg, in single or tandem form, along with (LAF-1 RGG)_2_-GFP. D. Plot showing variation in EI with B’s homotypic interaction at a constant heterotypic interaction (0.85) for B’s chain length of 20, 40 and 60 shown in colored lines. The top right plot shows enhanced EI for longer chain-length at 0.60 B’s homotypic interaction. The bottom right plot shows that for longer chain length, EI of 1.60 is achieved at lower B’s homotypic interactions. E. Position and segregation of charge as well as sequence charge decoration (SCD) of FUS LC variants and LAF-1 RGG shuffle variants. F. Segregation of charge on LAF-1 RGG, known to increase homotypic interactions^19^, partially blocks partitioning of FUS LC and its mutants. n ≥ 70 condensates per column. G. Plot showing variation of EI with heterotypic interaction for different constant homotypic interaction of scaffold shown in colored lines. Here the homotypic interaction of client is fixed at 0.85. The interaction strengths are normalized by homotypic interaction of scaffold molecule. The top right plot shows that at very high heterotypic interaction - greater than homotypic interaction of client - enrichment decreases with increasing homotypic interaction of scaffold. The bottom right plot shows that at constant EI, higher heterotypic interaction is required to overcome the strong homotypic interaction in scaffold for co-partitioning. Error bars represent 95% confidence intervals. Significance was calculated by one-way analysis of variance (ANOVA); ns P > 0.05, * P ≤ 0.05, ** P ≤ 0.01, *** P ≤ 0.001, and **** P ≤ 0.0001. Scale bar: 5 μm.

### Charge segregation on both IDR scaffolds

Having observed the impact of adding and segregating charge in FUS LC on its co-partitioning with LAF-1 RGG, we wondered to what extent segregation of charge on the other IDR, LAF-1 RGG, could also enhance enrichment of FUS LC variants due to strengthened heterotypic and homotypic interactions of the scaffold protein. In previous work on LAF-1 RGG, we shuffled its sequence (‘LAF-1 RGG shuffle B’) to segregate like charges, reducing SCD without altering residue composition, demonstrating that charge blockiness greatly increased its LLPS in vitro^20^. To determine its impact on co-partitioning with FUS LC, we generated strains to co-express (RGG shuffle B)_2_-GFP with FUS LC wildtype and charge mutants (Fig. 4E). Surprisingly, in all cases, condensates of RGG shuffle B showed reduced rather than enhanced partitioning of each FUS LC variant. In the case of Charge+42.seg, we found a statistically significant drop in its enrichment within LAF-1 RGG shuffle B condensates compared to LAF-1 RGG wildtype condensates (Fig. 4F). Although somewhat perplexing, computational modeling shed light on the likely presence of distinct regimes in which the partitioning behavior of two polymers can change drastically. At low or moderate homotypic interaction strength for each distinct polymer, a lowering of SCD for one polymer will increase its heterotypic interactions and ability to enrich the second polymer. Conversely, at high homotypic interaction strengths for each polymer, a lowered SCD would actually have the opposite effect, causing reduced enrichment of the second polymer (Fig. 4G). RGG shuffle B has greatly increased homotypic interactions^20^ that outcompete heterotypic interactions with FUS LC mutants, thereby reducing their enrichment in the condensed phase compared to LAF-1 RGG wildtype. Stated another way, once the homotypic interaction of one polymer (λ_AA_) reaches a critical value, the EI of the second polymer would drop as the first polymer’s self-interactions outcompete cross-polymer interactions.

Together, these results highlight how polypeptide chain features and competition between homotypic and heterotypic protein interactions can shape the propensity of one IDR to co-partition in the condensed phase of another. Using these basic rules, LLPS specificity can be enhanced through sequence evolution or engineering by shortening IDR chains or strengthening homotypic interactions. Future work will further define parameter space on the road to highly predictive models of IDR LLPS.

To provide a workflow for identifying polypeptide sequences capable of orthogonal condensation that may be useful for generating synthetic compartments, we have summarized the series of steps we used to assess LLPS specificity and then break it (Fig. 5A). To begin, IDPs must first be stripped of excess interaction domains and limited to an IDR region to unmask specificity (see Fig. 1C,D). Many IDR regions have high C_sat_s and are unable to condense under normal cellular expression levels. They must be modified to lower C_sat_, which can be achieved in two ways as demonstrated in this manuscript: multimerizing the sequence or adding ‘sticky’ residues like Tyr or Phe. Once an IDR is able to condense, LLPS specificity can be evaluated between two scaffolds. In our case of LAF-1 RGG and FUS LC, we found orthogonality in both of these C_sat_-lowering methods – (FUS LC wt)_3_ and +30Tyr – but this is not necessarily true of any two IDR sequences. After identifying these constructs as orthogonal, we then move to break this specificity through mutagenesis of one IDR. In the case of our two model IDRs, this was achieved by adding segregated charge to FUS LC, as demonstrated by (Charge+32.seg)_2_. Using this toolkit, IDR specificity can be evaluated and provide a foundation to engineering microcompartments within the cell.

**Fig. 5.**
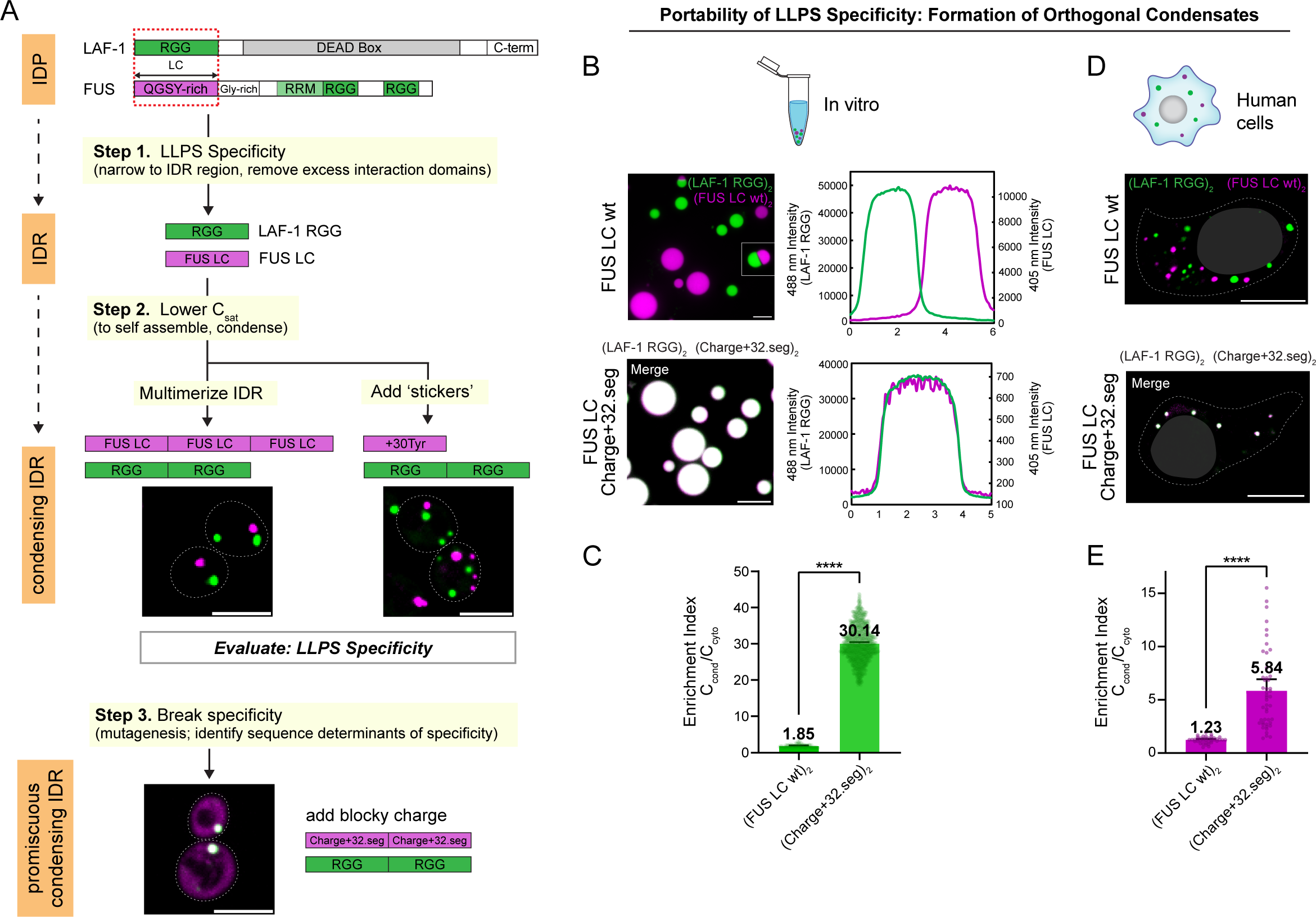
Rules for LLPS specificity are applicable broadly, in multiple biochemical and biological contexts. A. Workflow summarizing how to identify disordered proteins capable of orthogonal LLPS and then break specificity, using simple eukaryotic model cell to generate multiple distinct synthetic condensates. Step 1 induces specificity of IDPs by stripping away interaction domains that may promote co-assembly; leaving only IDRs. Step 2 lower C_sat_ to enable formation of condensates, via one of two methods: multimerization or addition of ‘sticky’ residues. Once two IDRs can condense, assess LLPS specificity through co-expression in cells. Once orthogonal sequences are identified, a third step may be taken to then break this specificity through mutagenesis, which further reveals the sequence determinants of LLPS specificity. B. Left: Biochemical reconstitution in vitro using purified proteins at concentrations above their C_sat_ to generate droplets. Top row: images showing formation of discrete condensates of (LAF-1 RGG)_2_-GFP and (FUS LC)_2_-BFP; right line scan. Bottom row: (LAF-1 RGG)_2_-GFP strongly co-partitions with FUS LC Charge+32.seg condensates; (FUS LC)_2_-BFP tracer; “merge” in white shows overlay of (LAF-1 RGG)_2_-GFP and (FUS LC)_2_-BFP tracer. Right: plot showing with line scan. Scale bar: 5 μm C. quantitation of micrographs from (B) showing enrichment of (LAF-1 RGG)_2_-GFP in FUS LC variant condensates. n ≥ 97 condensates per column. D. Selectivity of scaffold LLPS to form discrete condensates when expressed via transfection of human HEK293T cells. Top row: co-expression of (LAF-1 RGG)_2_-GFP and (FUS LC wt)_2_-mCherry leads to formation of discrete condensates. Lower: (LAF-1 RGG)_2_-GFP strongly co-partitioning with (FUS LC Charge+32.seg)_2_-mCherry when co-expressed in human HEK293T cells. Images are max projections. E. Quantitation of micrographs from (D) showing (FUS LC variant)_2_-mCherry enrichment in (LAF-1 RGG)_2_-GFP condensates. n ≥ 39 condensates per column. Error bars represent 95% confidence intervals. Significance calculated by a Welch’s t-test; ns P > 0.05, * P ≤ 0.05, ** P ≤ 0.01, *** P ≤ 0.001, and **** P ≤ 0.0001. Scale bar: 10 μm.

Next, we wanted to assess the extent to which the IDR polypeptide sequences and rules for LLPS specificity were portable to other biological and biochemical contexts. While our results seem to yield general principles in *S. cerevisiae*, we wanted to determine their applicability in minimal buffer, rather than complex cytosol, such as would be used to generate protocells and separately to determine the feasibility of generating orthogonal synthetic compartments in human cells for applications in cellular engineering. In short, we wanted to test the generalizability of our LLPS rules. To start, we biochemically reconstituted LAF-1 RGG and FUS LC condensates in vitro both at concentrations above their C_sat_. First, we assessed LLPS specificity between (LAF-1 RGG)_2_-GFP and (FUS LC wt)_2_-BFP condensates and demonstrated that they form orthogonal condensed phases that do not co-partition with one another (Fig. 5B). These scaffold-scaffold experiments confirm our results for scaffold-client schemes in vitro in Extended Data Fig. S2D-G and demonstrate that it is feasible to form discrete condensed phases in vitro in conditions that can be used to generate minimal cells. Next, we tested reconstitution of LAF-1 RGG and FUS LC Charge+32.seg mutant, we observed near complete overlap of signal (Fig. 5C, line scans to right, Extended Data Fig. S5A). This result indicates that loss of LLPS specificity in the FUS LC mutant Charge+32.seg is indeed specific to its altered sequence and not dependent on interactions with additional factors in cytosol such as protein or RNA. Next, we wanted to assess LLPS specificity rules in mammalian cells, an environment containing native FUS interaction partners that could interfere with assembly specificity. By co-transfecting HEK293T cells, both IDR constructs were expressed in tandem form to generate condensates. Cells co-expressing wildtype (LAF-1 RGG)_2_-GFP and (FUS LC wt)_2_-mCherry formed orthogonal condensates, with no overlapping signal (Fig. 5D). However, when FUS LC mutant (Charge+32.seg_2_-mCherry) was co-expressed with (LAF-1 RGG)_2_-GFP, we observed near complete overlap of signal as with the in vitro experiments, with strong enrichment of the FUS LC mutant in LAF-1 RGG condensates (Fig 5E, Extended Data Fig. S5B). The specific formation of orthogonal FUS LC condensates in human tissue culture and the loss of LLPS selectivity in the Charge+32.seg mutant provide further evidence that these chemical rules for LLPS specificity may be universal, independent of biological setting.

## Discussion

In this study, we sought to determine whether and how intrinsically disordered sequences contribute to specificity in the formation of biomolecular condensates. We set out to identify orthogonal IDR pairs that form discrete condensed phases and identified that FUS LC and LAF-1 RGG form spatially discrete compartments when co-expressed in cells. Using these two IDRs as a case study, we determined that LLPS specificity is encoded directly in the protein sequence: they are able to form discrete condensed phases in vitro in the absence of additional macromolecules such as RNA; additionally, this specificity is sufficient to enforce selectivity of condensation in the complex milieu of the cytoplasm. Further, we sought to determine the chemical principles governing LLPS specificity of these model IDPs to determine how selectivity can evolve or be engineered. We identified amino acids that contribute to the selectivity of FUS LC condensation and, conversely, that can be mutated to promote its promiscuous co-assembly with other condensates such as those formed by LAF-1 RGG. Additionally, we determined the contribution of polypeptide chain properties to LLPS specificity. We found that the extent of charge partitioning can have opposing effects, in some cases enhancing heterotypic interactions and in other cases causing greater increases in homotypic interaction strength, the combination of which determined IDR LLPS specificity.

A central biochemical question is which molecular interactions dictate the partitioning of IDPs to distinct subcellular complexes. There are likely multiple mechanisms. For some scaffold proteins that are capable of forming condensates on their own, selective assembly may be governed by the presence of folded domains that control specific interactions with RNA or client proteins^14, 16, 70^. Intriguingly, the presence of these tertiary molecules such as RNA can influence the co-mixing or co-partitioning of two IDRs^17, 71^. The similarity of IDRs can drive co-localization to a condensed phase; for example, the PLDs on transcription factors mediate co-partitioning to mediator condensates^72^. However, a code has yet to emerge that dictates the formation of multiple discrete phases of transcriptional condensates in the presence of orthogonal PLDs. A gap in knowledge has been understanding whether IDR sequences alone are sufficient to direct condensation into multiple separate membraneless compartments in a cell. To our knowledge, our study represents one of the first to characterize a pair of IDRs that are capable of forming spatially distinct condensates both in vitro and in cells and to reveal the sequence determinants that enforce the specificity of condensation.

An alternative hypothesis for the promiscuity of our FUS LC charge mutants could be a shift in material properties away from wildtype that allows for increased enrichment in the LAF-1 RGG condensed phase rather than increased heterotypic interactions of the two IDR sequences. To assess the biophysical properties of the condensed phase of FUS LC and its mutants Charge+32.seg and Charge+42.seg, we multimerized them to decrease C_sat_, enabling formation of condensed phases in yeast whose dynamics or liquidity could be characterized by fluorescence recovery after photobleaching (FRAP). We found that condensates for FUS LC wildtype and mutants all appeared to have very similar dynamics or half-time of recovery as well as similar mobile fraction (Extended Data Fig. S5C). This similarity in biophysical properties of FUS LC wildtype and mutant condensates indicates that the increased enrichment of FUS LC mutants in LAF-1 RGG condensates is unlikely related to a shift in its dynamics.

In considering how to build multiple discrete (non-mixing) membraneless compartments in cells, our study provides a number of design paradigms. It is feasible to start a design from relatively short non-mixing IDRs, using multimerization to lower C_sat_ to achieve formation of two or more discrete condensed phases in the cytosol of cells (Fig. 5A). Strong bias of amino acid composition may be sufficient to generate multiple non-mixing phases, as evidenced by use of an IDR such as FUS LC that is largely devoid of charged residues. It will be intriguing in future studies to test IDRs rich in hydrophobic residues and assess whether they can be used to form a third discrete condensed phase. Second, for IDRs that are relatively similar in composition of charged residues, their organization along the polypeptide chain can alter the balance of homotypic and heterotypic interactions. For example, we found that FUS LC mutant Charge+32 did not co-partition in LAF-1 RGG condensates, but FUS LC Charge+32.seg did (Fig. 2C, D). Further creating patchy charge on LAF-1 RGG shuffle B creates stronger homotypic interactions for itself which in turn diminish heterotypic co-mixing of FUS LC+32.seg (Fig. 4E, F). Similarly, charge blockiness influences association of transcription modulators into mediator condensates^72^. We propose it is necessary to consider not just the effects of mutation on heterotypic interactions, because the effects on homotypic interactions may be strong. Case in point, although Tyr residues should promote interaction with LAF-1 RGG^20^, their further addition to FUS LC was not adequate to cause co-partitioning in the RGG condensed phase. Instead, the additional Tyr lowered C_sat_, indicating greater homotypic interaction. Because the experimental impacts of changes to amino acid sequence on LLPS specificity are difficult to guess, computational modeling of homotypic and heterotypic IDR interactions greatly aids in deciphering these potential effects. Though recent studies have computationally shown formation of multilayered condensates^73^, our minimalistic model captures various scenarios like two distinct condensed phases, a co-existing phase, and a multi-layered phase that provide insight into the LLPS specificity of FUS LC and LAF-1 RGG. The simulation results emphasize how the interplay between homotypic and heterotypic interactions, serving as the two control parameters in the minimalistic model, determines whether co-mixed or discrete condensed phases form. We also emphasize that it may be challenging to target client proteins to a condensed phase by simply tagging with a similar IDR sequence^74^. A limitation of generating specificity in subcellular organization based exclusively on IDRs is that there exists an inherent competition between homotypic and heterotypic interactions. Alternatively, the addition of folded protein-protein interaction domains may be a more direct route to heterotypic assembly of components into mesoscale complexes. Indeed, addition of coiled-coils to an IDR scaffold and client has emerged as a highly efficient strategy to achieve targeting of an enzyme to a synthetic condensed phase^75^.

Overall, the findings of this study provide sequence determinants and conceptual paradigms that advance our understanding of how IDRs can drive selective partitioning into spatially distinct cytosolic condensates. These principles will aid the design of new sequences that are orthogonal to one another to form discrete condensed phases for construction of synthetic organelles to engineer pathways and behaviors of cells. Additionally, our work reinforces the perspective that to understand IDP subcellular partitioning, it is necessary to consider the chain level properties of the disordered sequence and how homotypic self-assembly may prevent heterotypic interactions with other IDRs.

## Methods

### Cloning

All plasmids were constructed using In-Fusion (Takara Bio) cloning and verified via Sanger sequencing (Azenta Life Sciences). LAF-1 RGG and FUS LC mutants were optimized for *Saccharomyces cerevisiae* and ordered as gBlocks from IDT and Twist Bioscience. Inserts were amplified by PCR from gBlocks and previously cloned plasmids. A yeast plasmid expressing the RGR scaffold was encoded in the integrating YIplac211 (URA3, ampicillin resistance (AmpR)) plasmid backbone downstream of the inducible GAL1 promoter as a tandem separated by meGFP. Yeast plasmids expressing FUS LC clients were encoded in the integrating YIplac128 (LEU2, ampicillin resistance (AmpR)) plasmid backbone downstream of the inducible GAL1 promoter and upstream of mScarlet and a triple PKI nuclear exit signal (NES). For higher valency constructs, the second version of LAF-1 RGG or FUS LC was inserted directly after the fluorescent tag and third versions (when applicable) directly after the second, all before the NES. For mammalian cell constructs, FUS LC wildtype and mutants were cloned into a pCDNA3.1 backbone.

### Yeast procedures

Standard methodologies were followed for all experiments involving *S. cerevisiae* ^76, 77^. The expressed proteins were constructed using a galactose-inducible GAL1 promoter and incorporated into YEF473 yeast URA3 and LEU2 loci using integrating vectors (YIplac211, YIplac128, respectively) cut with PpuMI or EcoRV and a standard lithium acetate transformation. All yeast strains used in this study are listed in Supplementary Table 1 and plasmids are listed in Supplementary Table 2.

To induce protein expression, yeast cells were first grown to saturation overnight in liquid YPD medium in a 25°C shaking incubator. Cell cultures were pelleted by centrifugation at 3000 rpm in a Sorvall Legend RT centrifuge (Thermo Scientific) and washed three times in sterile water. Cells were then diluted in YP + 2% raffinose and incubated in a 25°C shaking incubator for 6 to 8 hours. Finally, yeast cells were diluted to an optical density of 600 nm (OD_600_) of 0.3 in YP + 2% galactose, and induction proceeded overnight in the same shaking incubator. The final OD_600_ values of cultures used for experiments were between 0.4 and 0.8 at approximately 20 hours of galactose induction. For constructs involving tandem clients ((FUS LC wt)_2_, (Charge+32.seg)_2_), YP media with 0.1% galactose was used to limit overexpression and aggregation. For FRAP yeast colonies ((FUS LC wt)_3_, (Charge+32.seg)_2_, (Charge+42.seg)_2_), colonies were induced with 2% galactose approximately 8 hours prior to imaging and prepared as above.

### Protein Purification

Codon optimized LAF-1 RGG and FUS LC constructs were cloned into pET and pMAL expression vectors, respectively, by In-Fusion ligation (Takara Bio) and transformed into Rosetta 2 *E. coli* cells (Sigma Aldrich). For RGG constructs (RGG_2_ and RGG_2_-GFP), bacterial cultures were grown in Luria Broth (LB) supplemented with kanamycin and chloramphenicol at 37°C to an OD_600_ of 0.6–0.8 and expression was induced by 0.5 mM isopropyl β-d-1-thiogalactopyranoside (IPTG) at 16°C overnight. Cell cultures were centrifuged at 5,000 rpm in a Sorvall RC6+ centrifuge (Thermo Scientific) and the resulting pellets were collected and stored at −80°C. Bacterial pellets were thawed, then resuspended in lysis buffer (50 mM Tris-HCl, pH 7.5, 1 M NaCl, 20 mM imidazole, 1 mM β-mercaptoethanol) and complete EDTA-free protease inhibitor cocktail (Roche; Mannheim, Germany). Cells were lysed with 2 minutes of sonication at 50% power using a Branson Sonifier. Lysates were clarified by centrifugation at 13,000 rpm for 20 min in a Sorvall RC6+ centrifuge using a F21S-8×50y rotor (Thermo Fisher Scientific) at 37°C and incubated with 0.5 mL of Ni–NTA beads (Thermo Fisher Scientific) while rotating at room temperature for 1 hour. Beads were then washed three times with 10 mL of lysis buffer containing 40 mM imidazole. Proteins were eluted by lysis buffer containing 500 mM imidazole and 1 mM DTT. Eluted proteins were diluted to 3 mg/mL in lysis buffer containing 1 mM DTT and dialyzed overnight into storage buffer (20 mM Tris-HCl, pH 7.5, 500 mM NaCl, 1 mM DTT) using Slide-A-Lyzer membrane cassette (Thermo Fisher Scientific) with a 10 kDa size cutoff at room temperature. Proteins were concentrated by centrifugation in 4 mL of Amicon filter concentrators with a 10 kDa cutoff (Millipore Sigma; Burlington, MA). TCEP (1 mM) was added prior to snap freezing and storage at −80°C.

Constructs of FUS LC (FUS LC wildtype, GFP-FUS LC, and mutants) with an N-terminal MBP and C-terminal 6His tag were purified as stated above except that ampicillin was used instead of kanamycin for bacterial cell growth and expression and all steps of the purification were done at 4°C. Purification utilized only the C-terminal 6His tags on all FUS LC constructs. The N-terminal MBP tag was utilized to keep proteins soluble until experiments. A TEV cleavage site is present just upstream of the FUS LC sequence in construct and is cleaved prior to in vitro experiments. Tandem sequences of FUS LC wildtype with a BFP tag were cloned into a pMAL plasmid with an N-terminal MBP tag, H3C protease cleavage site, and a C-terminal 6His tag. This plasmid was transformed into Rosetta competent bacteria, expressed, and purified as described for FUS LC wildtype and the other described sequence variants and stored in identical conditions.

### Mammalian cell procedures

HEK293T cells (ATCC) were cultured in Eagle’s minimal essential medium (EMEM, Quality Biological) supplemented with 10% fetal bovine serum (Gibco™), 2 mM L-glutamine (Gibco™), and 10 U/mL penicillin-streptomycin (Gibco™) and maintained at 37°C in a humidified atmosphere with 5% CO_2_. Cells were split in 1:3 ratio every 3 days and had been passaged for less than 2 months, were negative for known infection, and experiments were done with confirmed viability >95% by Trypan Blue staining (Gibco™).

For expression of scaffolds, cells were seeded on a 24 well glass-bottom plate (Greiner Bio-One) at 70% confluency. 24 hours later, cells were transfected with 500 ng of each plasmid via lipofectamine 2000 (Invitrogen) according to manufacturer’s protocol.

### Microscopy

Images were collected on an Olympus IX81 inverted confocal microscope (Olympus Life Science) equipped with a Yokogawa CSU-X1 spinning disk, mercury lamp, 488- and 561-nm laser launches, and an iXon3 EMCCD camera (Andor). Multidimensional acquisition was controlled by MetaMorph software (Molecular Devices). Samples were illuminated using a 488-nm and/or a 561-nm laser and were imaged through a 100x, 1.4-NA oil immersion objective. Z stacks were collected at 0.3 μm steps. Yeast samples were placed in custom-fabricated acrylic gasket chambers adhered to glass coverslips (#1.5 glass thickness; Corning Inc.) treated with concanavalin A (ConA, 2 mg/mL, rinsed after drying overnight) for immobilization and immediately imaged.

For yeast fluorescence recovery after photobleaching (FRAP) experiments, a Roper iLas2 photoactivation system controlling a 405-nm laser was used. Individual condensates were selected and photobleached, and fluorescence recovery in the bleached region was analyzed in ImageJ after using histogram matching bleach correction to correct against photobleaching due to long period of laser exposure. Z stacks were collected to visualize the condensates at the 561-nm wavelength using a 100x, 1.4-NA oil immersion objective. Imaging of condensates in vitro was performed on the same microscope as above, except in the case of experiments in Fig 5. Proteins were thawed at 50°C and scaffold proteins (tandem LAF-1 RGG, FUS LC, and FUS LC mutant constructs) were diluted to the desired concentration in buffer containing 150 mM NaCl and 20 mM Tris-HCl, pH 7.5, 0.5-2.5 μg TEV protease (depending on FUS LC concentration), and 1 mM DTT in a 20 μL volume. This mixture was incubated at 37°C for 30 min to cleave off the MBP tag and then placed into the same custom-fabricated acrylic gasket chambers adhered to glass coverslips (#1.5 glass thickness; Corning Inc.), which were passivated with 10 mg/mL BSA. Condensate formation was allowed to proceed for 20 minutes at ambient temperature (approximately 22°C) before imaging. Client proteins (RGG_2_-GFP and GFP-FUS LC) were added to scaffold mixtures at a final concentration of 0.2 μM.

For experiments in which both FUS LC and LAF-1 RGG were used at concentrations above their C_sat_, (Fig 5B, C) the droplets were imaged on a CrestOptics X-Light V3 spinning disk confocal microscope equipped with 405 nm, 445 nm, 488 nm, 520 nm, 555 nm, and 630 nm laser launches, pco.edge sCMOS cameras, and an Okolab enclosure. Image acquisition controlled by VisiView software. Aliquots of purified RGG-GFP-RGG, MBP-FUS LC-BFP-FUS LC, and MBP-FUS LC Charge+32.seg were thawed at 50°C. FUS LC Charge+32.seg polypeptides were cleaved from N-terminal MBP tags with TEV protease for 20 minutes at ambient temperature in pre-passivated imaging wells as described above. FUS LC-BFP-FUS LC were cleaved away from the N-terminal MBP tag using H3C protease at 42°C for 10 minutes. Following cleavage and droplet formation of 8 µM FUS LC Charge+32.seg, 8 µM RGG-GFP-RGG and 0.2 µM FUS LC-BFP-FUS LC were added, and the mixture was incubated at ambient temperature for 10 minutes before imaging. Final buffer conditions: 20 mM Tris-HCl, pH7.5, 250 mM NaCl, and 1 mM DTT. To evaluate specificity of RGG and FUS LC directly, RGG-GFP-RGG and FUS LC-BFP-FUS LC were prepared as above and added to a pre-passivated imaging well to a final concentration of 2 µM each in buffer containing 20 mM Tris-HCl, pH7.5, 100 mM NaCl, and 1 mM DTT.

### Turbidity Measurements

Proteins were thawed at 50°C as above and diluted to 10 μM in 150 mM NaCl and 20 mM Tris-HCl, pH 7.5, and 1 mM DTT in a 60 μL volume. Protein mixtures were placed in pre-warmed quartz cuvettes with a 10 mm path length (Starna Cells, Inc). Cuvettes were then inserted into a preheated (50°C) Cary 3500 UV−Vis spectrophotometer controlled by an Agilent multizone Peltier temperature controller (Agilent Technologies). The samples were cooled to 1°C at a rate of 1°C/min while measuring the absorbance at 600 nm.

### Image Analysis

Image segmentation and analysis was performed using custom-written macros in ImageJ to partially automate the process and enable collection of larger sample sizes. A multistep process was employed to determine enrichment indices with a series of checkpoints throughout.

First, z stacks (21 slices, step size 0.3 μm) were converted into MAX projections to visualize all condensates in a single plane. Next, each image was manually thresholded to create a mask such that each cell could be easily found via the “Analyze Particles” function and individually cropped into a series of tif files with one cell per. Since each cell had varying expression levels, analysis had to be done on a cell-by-cell basis for more accurate cytoplasmic and subsequent enrichment values. During the selection of cells for analysis, cells with significantly different expression in either channel were excluded as well as any cells that showed considerable travel of condensates through the stack (most notably a linear smear in the MAX projection). Once individual cell files were created, a script thresholded each image in the 561 nm channel according to an input value predetermined by the user. This threshold was determined by multiplying the background of the image by a series of integers and referring to the original MAX projection 561 channel for the best fit of visualizing cytoplasm. Thresholds varied from 2x to 10x background depending on protein expression, imaging fluctuations, etc. Once a threshold was determined, it was used in the 561 nm channel to create a mask in each cell, create an ROI, and measure area, mean, standard deviation, mode, minimum, maximum, and integrated density of the region in both the 488 nm and 561 nm channels. These values will be referred to as “initial cytoplasm measurements” as they include condensate values that will inflate mean cytoplasm values and provide an inaccurate denominator for enrichment indices. Next, thresholds were calculated to find and remove RGR condensates in the 488 nm channel by adding the 488 nm channel initial cytoplasm mode to the standard deviation. The resulting masks and ROIs in the 488 nm channel should only include pixel values in RGR condensates which are then subtracted from the initial cytoplasm ROIs through the XOR function. For condensing FUS LC mutant condensates, an extra step is added to also subtract condensates in the 561 nm channel. These new ROIs were then measured in both channels and results deemed “accurate cytoplasm measurements”. From here, a final threshold was calculated by using the 488 nm accurate cytoplasm mean and adding two standard deviations. The final script created ROIs for each condensate, excluding those smaller than 9 pixels in area. For each condensate, the macro found the maximum pixel value within the RGR condensate in the 488 nm channel and then measured that pixel in both channels. These are the values that were used in the numerator of the enrichment index calculation. For an alternative EI calculation method using the average 561 nm pixel value throughout the thresholded condensate, see Extended Data Table S1. With all necessary data extracted from the images, enrichment indices were calculated for each condensate in both channels. This was done by using background-corrected peak values from the last step (intensity_condensate_) divided by the background-corrected accurate cytoplasm mean (intensity_cytosol_). These values were compiled and filtered by eliminating condensate EI values outside of two standard deviations of the 488 nm channel mean.

Similarly, enrichment indices for GFP-tagged RGG constructs within in vitro condensates were calculated simply by dividing the peak condensate value by the image background.

For FUS LC mutants with additional Tyr residues (+20Tyr and +30Tyr), expression was incredibly low and quantification had to be done as follows. Cells were manually cropped from single slices (MAX projections could not be used due to coarseness of cytoplasm and abundance of condensates per cell) that showed both RGR condensates and mScarlet expression above background. The cytoplasmic values for each cell were generated by manually drawing an ROI around as much cytoplasm as possible without including condensates in either channel. Then, the maximum value of each RGR condensate was found (generally the middle pixel of the condensate) and recorded as the 488 nm peak value. Since the 561 nm cytoplasmic signal was so low and variable, a circle with a diameter of 3 pixels was drawn centered around the peak 488 nm pixel. The average of this small circle was used as the 561 nm peak value. Then, as in the semi-automated version of quantification, these peak and cytoplasmic values were background-corrected and used to calculate EI values per condensate and filtered as above.

Scripts used to generate the macros used for analysis in ImageJ are available upon request.

### Molecular Model and Simulation

The coarse-grained (CG) coexistence simulation protocol used in the study was based on the previously developed coarse-grained slab simulation for proteins^61, 78^. We considered two polymer chains, A and B, of length 20-mer each unless otherwise noted. The bonded interactions between the monomers were modeled using a harmonic potential as:

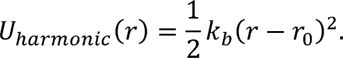

Where *k_b_* is the harmonic spring constant equal to 20 kcal/mol. Å and *r_0_* is the equilibrium bond length equal to 3.8 Å. The nonbonded interactions between the monomer are defined using a modified Lennard-Jones (LJ) potential^79^,

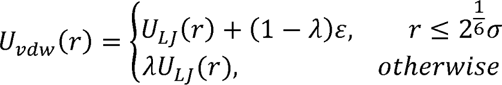

Where λ controls the attractive well-depth and ∪*_LJ_* is the Lennard-Jones potential

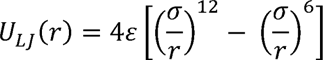

In our model, the LJ diameter (σ) of the monomers was set to 5 Å and the interaction parameter (ε) was set to 0.20 kcal/mol. The λ_AA_ is fixed as 1 between the monomers of polymer A, while it is varied from 0 to 1 for λ_BB_ (between monomers of B) and λ_AB_ (between A and B monomers).

The simulations were performed using the HOOMD-blue 2.9.3 simulation software package^80^ in a cuboid shape box with x and y dimensions of 100 Å and z dimensions of 750 Å and a timestep of 10 fs. The temperature was set at 300K using Langevin thermostat with residue friction coefficient, γ = m_i_/t_damp_, where m_i_ is mass of each monomer (100 g/mol) and t_damp_ is damping factor, which was set to 1000 ps. Simulations were performed using 250 chains each of polymer A and B. We conducted scanning simulations with a constant λ_AB_ while altering λ_BB_ from 0 to 1 in increments of 0.02, and vice versa. For each data point, we ran simulations for 100 ns. Additionally, we performed scans across straight lines both above and below the diagonal.

For studying the role of the client’s chain length, simulations were also performed with 40-mer and 60-mer B polymers, keeping the total number of monomers fixed to the same as for 20-mer. The enrichment index (EI) is computed as the ratio of B’s concentration inside A to B’s total solution concentration. Due to a small finite number of B chains in the simulation setup, we use solution concentration instead of dilute phase concentration to avoid numerical issues when B concentration in the dilute phase goes to 0.

### Statistics and Reproducibility

Experiments were reproducible. All statistical analyses were performed in GraphPad Prism 9. To test the significance of two categories, an unpaired Welch’s t-test was used. To test significance of more than two categories, a one-way ANOVA was used. In all cases, ns indicates not significant (P < 0.05), * P ≤ 0.05, ** P ≤ 0.01, *** P ≤ 0.001, and **** P ≤ 0.0001.

## Acknowledgements.

We thank the E. Bi and J. Shorter labs for sharing yeast strains and plasmids, A. Stout and the Penn CDB Microscopy Core for imaging and support. This study was supported by National Institute of Health grants, including National Institute of Biomedical Imaging and Bioengineering grant, EB028320 (M.C.G) and NIGMS R01GM136917 (J.M.). Additionally, the work was partly funded by National Science Foundation (NSF) MRSEC Seed grant, DMR1720530 (M.C.G) and Welch Foundation A-2113-20220331 (J.M.). Work by M.C.G., D.A.H., and M.G. supported in part by the U.S. Department of Energy (DOE), Office of Science, Basic Energy Sciences (BES), under Award # DE-SC0007063 (M.C.G).

## Author Contributions

M.C.G and J.M. conceptualized the project. M.C.G., R.M.W., and M.V.G. designed the experiments. E.G. performed initial IDP cloning and strain generation for yeast co-expression screening. B.X. generated clones and strains for initial characterization of FUS LC multimers. R.M.W. performed cloning, strain generation, and imaging for all yeast experiments following initial screenings, including FUS LC mutagenesis. K.A.S. designed predicted FUS LC mutants. M.V.G. purified recombinant FUS LC and mutants and performed in vitro characterization of LLPS. K.A.S. generated minimalistic polymer model for two-component phase separation aided by R.M.R.. R.M.W. and M.V.G. analyzed experimental data. W.W. transduced HEK293T cells and imaged constructs. M.G. and M.V.G. purified recombinant (FUS LC)_2_-BFP. M.C.G., R.M.W., K.A.S., M.V.G., and J.M. wrote the manuscript.

## Supplemental Information

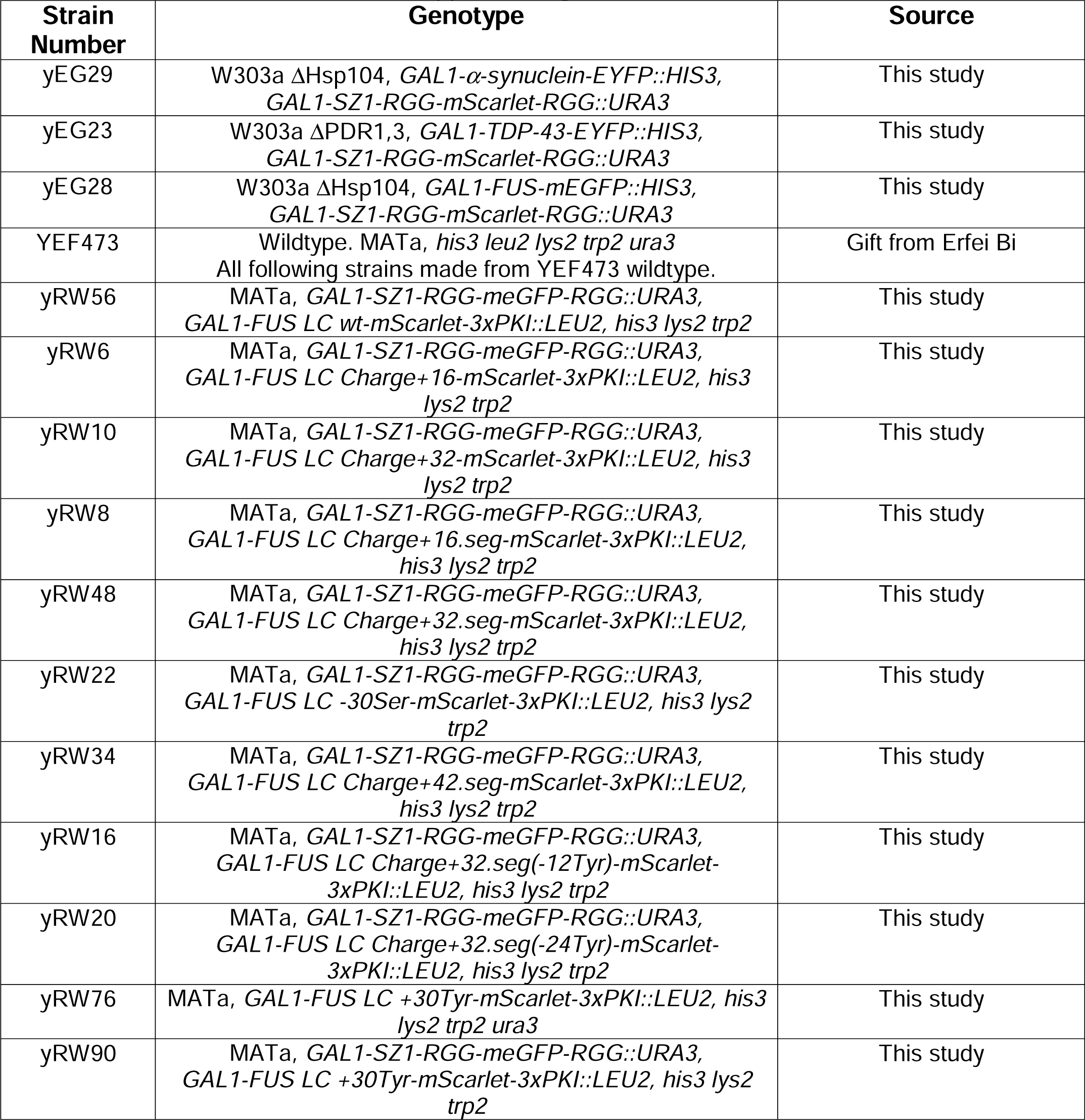
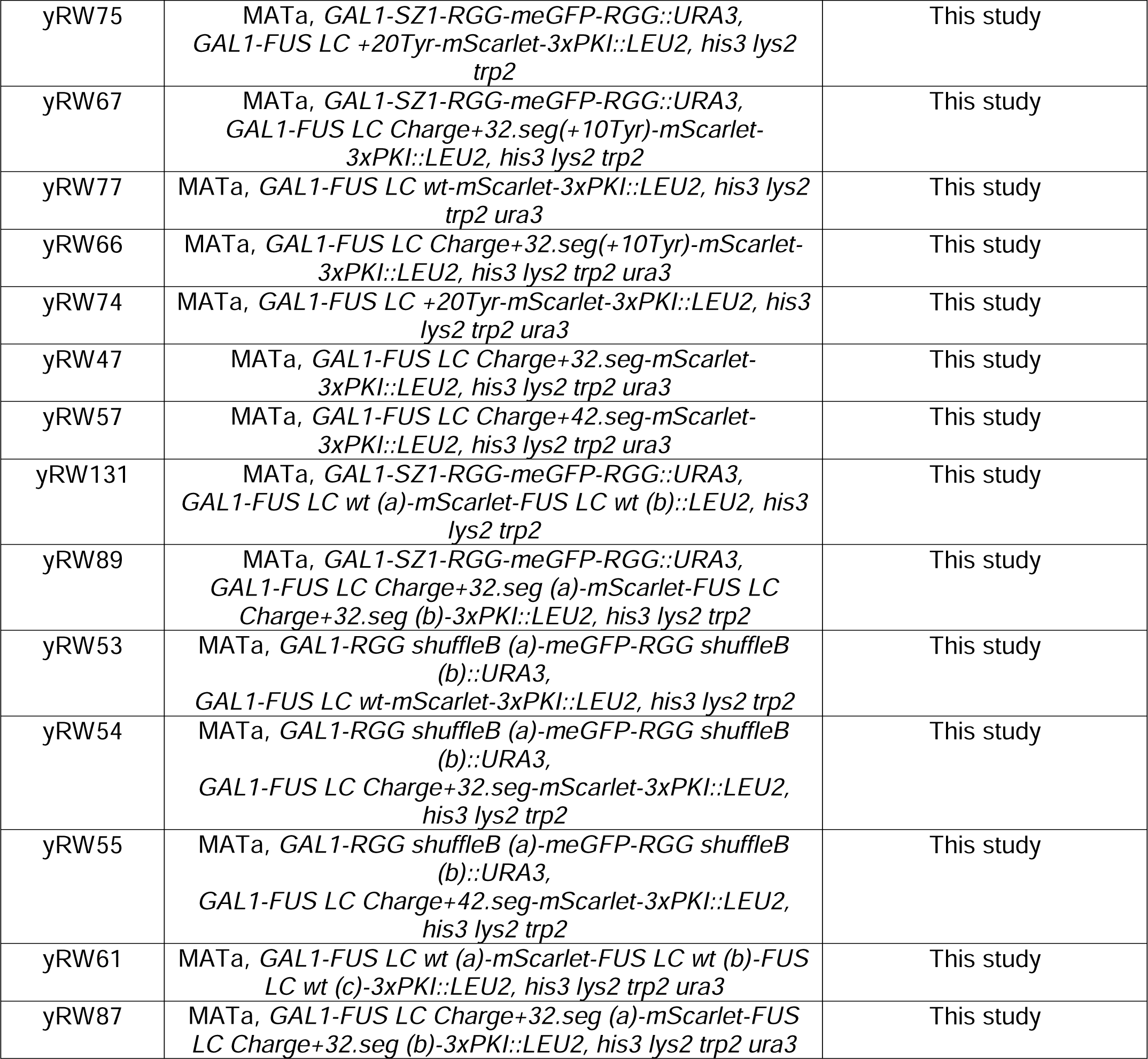
Supplementary Table 1.

Supplementary Table 1
| Strain Number | Genotype | Source |
| --- | --- | --- |
| yEG29 | W303a ΔHsp104, *GAL1-α-synuclein-EYFP::HIS3, GAL1-SZ1-RGG-mScarlet-RGG::URA3* | This study |
| yEG23 | W303a ΔPDR1,3, *GAL1-TDP-43-EYFP::HIS3, GAL1-SZ1-RGG-mScarlet-RGG::URA3* | This study |
| yEG28 | W303a ΔHsp104, *GAL1-FUS-mEGFP::HIS3, GAL1-SZ1-RGG-mScarlet-RGG::URA3* | This study |
| YEF473 | Wildtype. MATa, *his3 leu2 lys2 trp2 ura3*<br>All following strains made from YEF473 wildtype. | Gift from Erfei Bi |
| yRW56 | MATa, *GAL1-SZ1-RGG-meGFP-RGG::URA3, GAL1-FUS LC wt-mScarlet-3xPKI::LEU2, his3 lys2 trp2* | This study |
| yRW6 | MATa, *GAL1-SZ1-RGG-meGFP-RGG::URA3, GAL1-FUS LC Charge+16-mScarlet-3xPKI::LEU2, his3 lys2 trp2* | This study |
| yRW10 | MATa, *GAL1-SZ1-RGG-meGFP-RGG::URA3, GAL1-FUS LC Charge+32-mScarlet-3xPKI::LEU2, his3 lys2 trp2* | This study |
| yRW8 | MATa, *GAL1-SZ1-RGG-meGFP-RGG::URA3, GAL1-FUS LC Charge+16.seg-mScarlet-3xPKI::LEU2, his3 lys2 trp2* | This study |
| yRW48 | MATa, *GAL1-SZ1-RGG-meGFP-RGG::URA3, GAL1-FUS LC Charge+32.seg-mScarlet-3xPKI::LEU2, his3 lys2 trp2* | This study |
| yRW22 | MATa, *GAL1-SZ1-RGG-meGFP-RGG::URA3, GAL1-FUS LC -30Ser-mScarlet-3xPKI::LEU2, his3 lys2 trp2* | This study |
| yRW34 | MATa, *GAL1-SZ1-RGG-meGFP-RGG::URA3, GAL1-FUS LC Charge+42.seg-mScarlet-3xPKI::LEU2, his3 lys2 trp2* | This study |
| yRW16 | MATa, *GAL1-SZ1-RGG-meGFP-RGG::URA3, GAL1-FUS LC Charge+32.seg(-12Tyr)-mScarlet-3xPKI::LEU2, his3 lys2 trp2* | This study |
| yRW20 | MATa, *GAL1-SZ1-RGG-meGFP-RGG::URA3, GAL1-FUS LC Charge+32.seg(-24Tyr)-mScarlet-3xPKI::LEU2, his3 lys2 trp2* | This study |
| yRW76 | MATa, *GAL1-FUS LC +30Tyr-mScarlet-3xPKI::LEU2, his3 lys2 trp2 ura3* | This study |
| yRW90 | MATa, *GAL1-SZ1-RGG-meGFP-RGG::URA3, GAL1-FUS LC +30Tyr-mScarlet-3xPKI::LEU2, his3 lys2 trp2* | This study |
| yRW75 | MATa, GAL1-SZ1-RGG-meGFP-RGG::URA3,<br>GAL1-FUS LC +20Tyr-mScarlet-3xPKI::LEU2, his3 lys2<br>trp2 | This study |
| yRW67 | MATa, GAL1-SZ1-RGG-meGFP-RGG::URA3,<br>GAL1-FUS LC Charge+32.seg(+10Tyr)-mScarlet-<br>3xPKI::LEU2, his3 lys2 trp2 | This study |
| yRW77 | MATa, GAL1-FUS LC wt-mScarlet-3xPKI::LEU2, his3 lys2<br>trp2 ura3 | This study |
| yRW66 | MATa, GAL1-FUS LC Charge+32.seg(+10Tyr)-mScarlet-<br>3xPKI::LEU2, his3 lys2 trp2 ura3 | This study |
| yRW74 | MATa, GAL1-FUS LC +20Tyr-mScarlet-3xPKI::LEU2, his3<br>lys2 trp2 ura3 | This study |
| yRW47 | MATa, GAL1-FUS LC Charge+32.seg-mScarlet-<br>3xPKI::LEU2, his3 lys2 trp2 ura3 | This study |
| yRW57 | MATa, GAL1-FUS LC Charge+42.seg-mScarlet-<br>3xPKI::LEU2, his3 lys2 trp2 ura3 | This study |
| yRW131 | MATa, GAL1-SZ1-RGG-meGFP-RGG::URA3,<br>GAL1-FUS LC wt (a)-mScarlet-FUS LC wt (b)::LEU2, his3<br>lys2 trp2 | This study |
| yRW89 | MATa, GAL1-SZ1-RGG-meGFP-RGG::URA3,<br>GAL1-FUS LC Charge+32.seg (a)-mScarlet-FUS LC<br>Charge+32.seg (b)-3xPKI::LEU2, his3 lys2 trp2 | This study |
| yRW53 | MATa, GAL1-RGG shuffleB (a)-meGFP-RGG shuffleB<br>(b)::URA3,<br>GAL1-FUS LC wt-mScarlet-3xPKI::LEU2, his3 lys2 trp2 | This study |
| yRW54 | MATa, GAL1-RGG shuffleB (a)-meGFP-RGG shuffleB<br>(b)::URA3,<br>GAL1-FUS LC Charge+32.seg-mScarlet-3xPKI::LEU2,<br>his3 lys2 trp2 | This study |
| yRW55 | MATa, GAL1-RGG shuffleB (a)-meGFP-RGG shuffleB<br>(b)::URA3,<br>GAL1-FUS LC Charge+42.seg-mScarlet-3xPKI::LEU2,<br>his3 lys2 trp2 | This study |
| yRW61 | MATa, GAL1-FUS LC wt (a)-mScarlet-FUS LC wt (b)-FUS<br>LC wt (c)-3xPKI::LEU2, his3 lys2 trp2 ura3 | This study |
| yRW87 | MATa, GAL1-FUS LC Charge+32.seg (a)-mScarlet-FUS<br>LC Charge+32.seg (b)-3xPKI::LEU2, his3 lys2 trp2 ura3 | This study |

Supplementary Table 2.

Supplementary Table 2
| Number | Plasmid | Source |
| --- | --- | --- |
| pMV6 | YIplac211-proGAL1-SZ1-RGG-meGFP-RGG::URA3 | Garabedian et al.<br>2021 |
| pRW42 | YIplac211-proGAL1-SZ1-RGG-mScarlet-RGG::URA3 | This study |
| pRW38 | YIplac128-proGAL1-FUS LC wt-mScarlet-3xPKI::LEU2 | This study |
| pRW56 | YIplac128-proGAL1-FUS LC Charge+16-mScarlet-<br>3xPKI::LEU2 | This study |
| pRW6 | YIplac128-proGAL1-FUS LC Charge+32-mScarlet-<br>3xPKI::LEU2 | This study |
| pRW47 | YIplac128-proGAL1-FUS LC Charge+16.seg-mScarlet-<br>3xPKI::LEU2 | This study |
| pRW19 | YIplac128-proGAL1-FUS LC Charge+32.seg-mScarlet-<br>3xPKI::LEU2 | This study |
| pRW12 | Ylplac128-proGAL1-FUS LC -30Ser-mScarlet-3xPKI::LEU2 | This study |
| pRW29 | Ylplac128-proGAL1-FUS LC Charge+42.seg-mScarlet-3xPKI::LEU2 | This study |
| pRW9 | Ylplac128-proGAL1-FUS LC Charge+32.seg(-12Tyr)-mScarlet-3xPKI::LEU2 | This study |
| pRW11 | Ylplac128-proGAL1-FUS LC Charge+32.seg(-24Tyr)-mScarlet-3xPKI::LEU2 | This study |
| pRW44 | Ylplac128-proGAL1-FUS LC +20Tyr-mScarlet-3xPKI::LEU2 | This study |
| pRW45 | Ylplac128-proGAL1-FUS LC +30Tyr-mScarlet-3xPKI::LEU2 | This study |
| pRW34 | Ylplac128-proGAL1-FUS LC Charge+32.seg(+10Tyr)-mScarlet-3xPKI::LEU2 | This study |
| pRW72 | Ylplac128-proGAL1-FUS LC wt (a)-mScarlet-FUS LC wt (b)::LEU2 | This study |
| pRW46 | Ylplac128-proGAL1-FUS LC Charge+32.seg (a)-mScarlet-FUS LC Charge+32.seg (b)-3xPKI::LEU2 | This study |
| pRW27 | Ylplac211-proGAL-RGG shuffleB (a)-meGFP-RGG shuffleB (b)-3xPKI::URA3 | This study |
| pRW61 | Ylplac128-proGAL1-FUS LC wt (a)-mScarlet-FUS LC wt (b)-FUS LC wt (c)-3xPKI::LEU2 | This study |

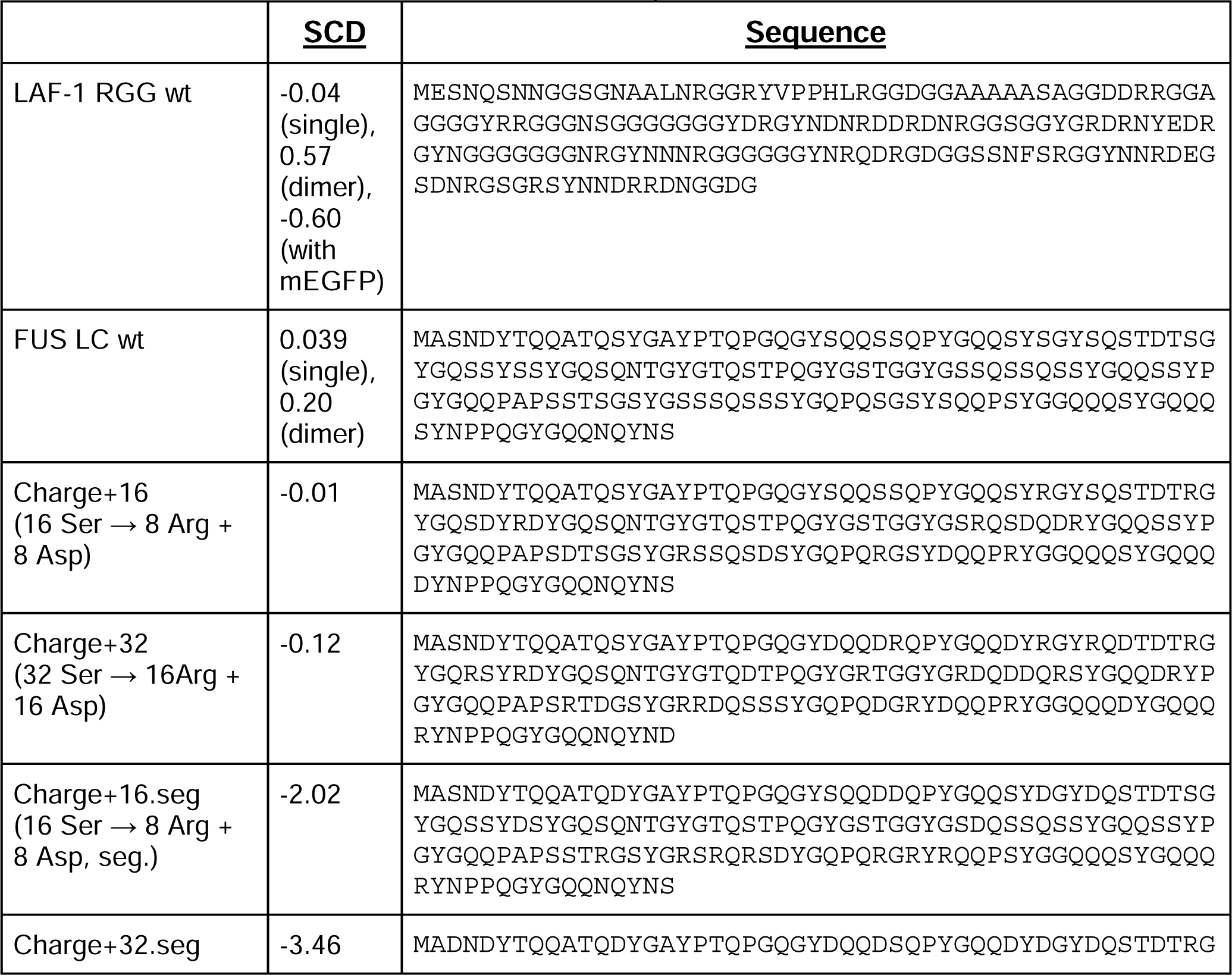
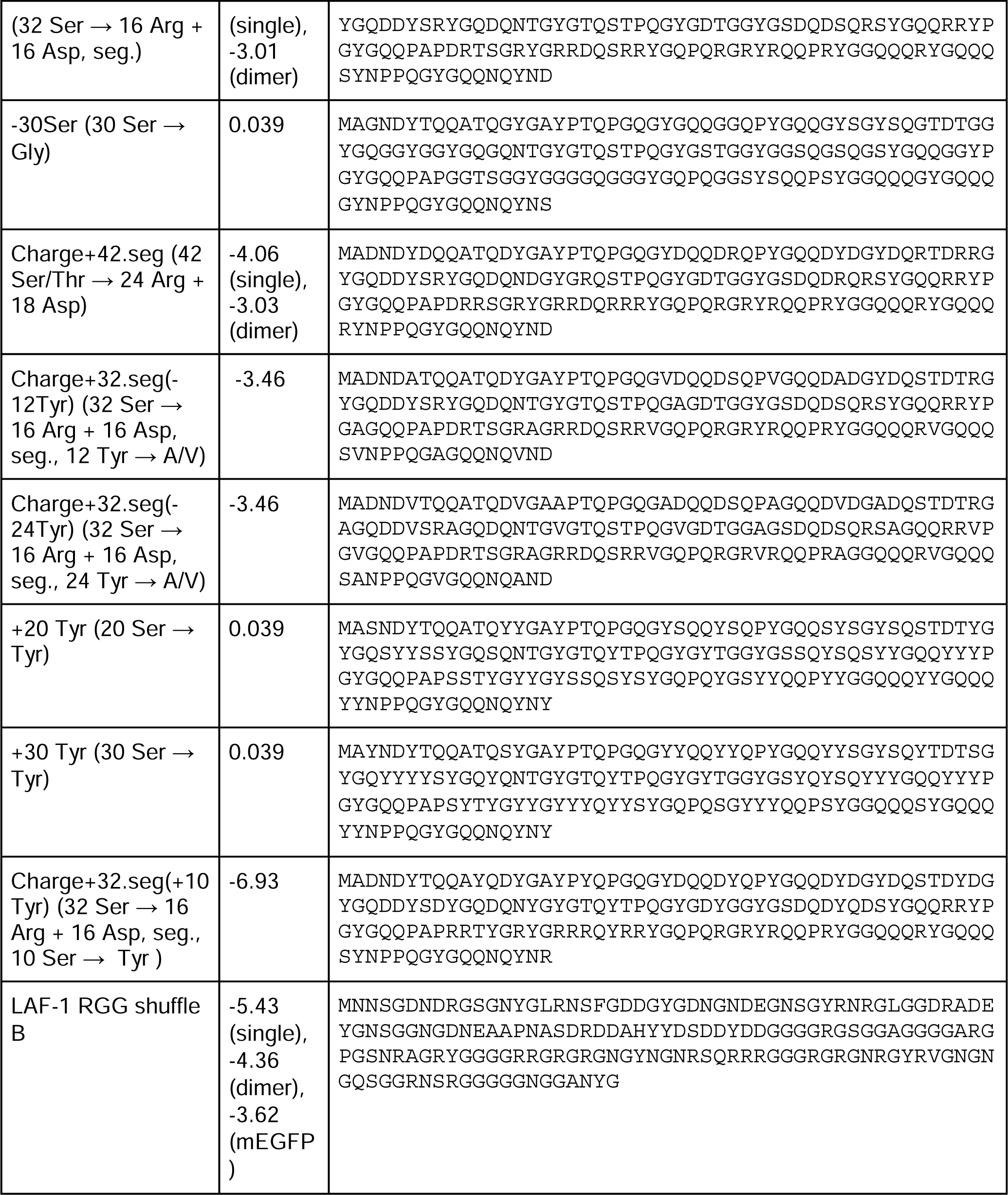
Protein Sequences.

## Figure Legends

**Extended Data Fig. S1. Supplemental data for Figure 1.** A. Quantitation of IDP enrichment in LAF-1 RGG condensates from IDP co-expression screening in yeast; images from Fig 1B. EI values for 3 IDPs in (LAF-1 RGG)_2_-mScarlet condensates. n ≥ 31 condensates per column. B. Plot showing possible range of enrichment indices in LAF-1 RGG condensates. Minimum value of 1. Maximum value based on LAF-1 RGG mScarlet (client) co-partitioning to (LAF-1 RGG)_2_-GFP condensates. n ≥ 37 condensates per column. C. Schematic of CG slab at 300K consisting of FUS LC (green) and LAF-1 RGG (magenta). Distinct phases are observed with FUS LC forming a condensed phase and LAF-1 RGG sticking at its interface. Significance was calculated by one-way analysis of variance (ANOVA); ns P > 0.05, * P ≤ 0.05, ** P ≤ 0.01, *** P ≤ 0.001, and **** P ≤ 0.0001. Data relevant to Figure 1.

**Extended Data Fig. S2. Additional Images and controls for FUS LC Mutant Analysis.** A. Non-mixing behavior of FUS LC wildtype persists in mutants Charge+16 and Charge+32 when adding balanced charge. B. Average numbers of (LAF-1 RGG)_2_-GFP condensates in co-expressing FUS LC mutant strains are within twofold of (LAF-1 RGG)_2_ and FUS LC wildtype strain. Light blue box indicates range within twofold of wildtype. n ≥ 30 cells per column noted below column mean. C. Expression of FUS LC mutant constructs within co-expressed strains, normalized to FUS LC wildtype expression on day of imaging. Light magenta box indicates range within twofold of wildtype. n ≥ 30 cells per column. Error bars represent 95% confidence intervals. Significance was calculated by one-way analysis of variance (ANOVA); ns P > 0.05, * P ≤ 0.05, ** P ≤ 0.01, *** P ≤ 0.001, and **** P ≤ 0.0001. Scale bar: 5 μm. Data relevant to Figure 2.

**Extended Data Fig. S2 (continued). LLPS specificity of purified FUS LC mutants and LAF-1 RGG proteins from biochemical reconstitution.** D. Partitioning of GFP-tagged (LAF-1 RGG)_2_ to FUS LC wildtype and condensates in vitro. E. Quantitation of enrichment of (LAF-1 RGG)_2_-GFP in condensed phase versus continuous phase from images in (D) 10 minutes after addition of (LAF-1 RGG)_2_-GFP mutants to pre-formed FUS LC condensates (50 μM of each FUS LC construct). n = 20 condensates per column. F. Brightfield images of wells containing various concentrations of FUS LC mutants, showing their phase boundaries. Estimated C_sat_ (right). G. Turbidity assays of FUS LC mutants (average of four independent trials) showing increased transition temperatures compared to wildtype. Error bars represent 95% confidence intervals. Scale bar: 10 μm. Data relevant to Figure 2.

**Extended Data Fig. S3. Additional information from polymer model simulations.** A. Density profiles of polymer B and EI plot for λ_AB_ = λ_BB_ case. B. Colormap of EI from CG scanning simulation with varying λ_AB_ and λ_BB_. C. Plot showing variation of EI as a function of λ_AB_ at λ_BB_ = 0 highlights the complex coacervate case. D. Plot showing variation of EI as a function of λ_BB_/λ_AB_ at different constant λ_AB_ values. Maximum enrichment is observed when ratio of λ_BB_/λ_AB_ is close to 1. E. Density profiles of polymer A and B corresponding to the cartoons in Fig. 3E. F. Density profile from single component CG slab for varying interaction strength. For interaction strength for 0.75 or above, chains start to condense. G. Simultaneous changes in homotypic and heterotypic interaction parameters can yield experimentally observed plateauing in EI. Plot shows variation of EI (right) along an arbitrarily defined change in λ_AB_ and λ_BB_ values (blue line shown in heat map (left)). Right plot shows plateauing of EI with increasing λ_BB_ (bottom x-axis) and λ_AB_ (upper x-axis). For +20Tyr and +30Tyr, increasing tyrosine will increase both homotypic and heterotypic interactions and scanning simulation shows one can obtain similar EI values as the FUS LC wildtype. For clarity, we only show one representative combination of λ_AB_ and λ_BB_, but there are several other possible combinations that can yield similar EI values as the FUS LC wildtype. Data relevant to Figure 3.

**Extended Data Fig. S4. Results from simulations varying polymer lengths.** Variation of EI with B’s homotypic interaction at 0.80 heterotypic interaction. At low heterotypic interaction (≤ 0.80) longer chain-length do not enhance co-partitioning. Data relevant to Figure 4.

**Extended Data Fig. S5. Additional images relevant to LLPS specificity and formation of orthogonal condensates in vitro and in mammalian cells.** A. Biochemical reconstitution in vitro using purified proteins at concentrations above their C_sat_, to generate droplets. (LAF-1 RGG)_2_-GFP strongly co-partitions with FUS LC Charge+32.seg condensates; (FUS LC)_2_-BFP tracer; “merge” in white shows overlay. B. Loss of LLPS specificity for FUS LC mutant charge+32.seg in HEK293T cells co-transfected with (LAF-1 RGG)_2_-GFP; (FUS LC Charge+32.seg)_2_-mCherry strongly co-partitions with (LAF-1 RGG)_2_-GFP condensates. Scale bar: 10 μm. C. FRAP plots from condensates of FUS LC constructs when expressed as multimers. Left: Triple version of FUS LC wildtype. Middle: Tandem version of Charge+32.seg. Right: Tandem version of Charge+42.seg. Plots show 95% confidence intervals. Samples sizes are shown on each plot. Data relevant to Figure 5.

**Extended Data Table S1. Enrichment indices of FUS LC clients using mean client values.** The second and third columns show the FUS LC client EIs as calculated throughout the manuscript using the max 488 nm pixel in each channel as the peak value in EI calculation (along with 95% confidence intervals). The fourth and fifth columns show values if EIs were instead calculated by averaging all 561 nm pixel values within the thresholded LAF-1 RGG condensate, frequently averaging in artificially low values due to variability in signal (also with 95% confidence intervals). Due to low expression, mutants +20Tyr and +30Tyr were analyzed using a different technique and could not be quantified with this alternative procedure (see Methods).

**Supplementary Fig. 1: Rigid polymer simulation.** A. Probability distribution of radius of gyration R_g_ from single chain simulations for different angle constant (kq). B. Cartoon representation of single chain at different rigidity. C. Variation in EI as a function of λ_BB_ at a constant λ_AB_ (0.85) for different angle constants. D. Cartoon representations of two component system with flexible polymer A (blue) and rigid polymer B (red). Cartoons from top to bottom represents the scenarios with increasing λ_BB_ at a constant λ_AB_.

