## Supporting Information for "Determinants of Disordered Protein Co-Assembly Into Discrete Condensed Phases"

[1] Department of Cell and Developmental Biology, Perelman School of Medicine, University of Pennsylvania, PA 19104
[2] Artie McFerrin Department of Chemical Engineering, Texas A&M University, TX 77843
[3] Present Address: Bioengineering Graduate Program, Rice University, Houston TX 77005
[4] Bioengineering Graduate Program, University of Pennsylvania, PA 19104
[5] Chemical and Biomolecular Engineering Graduate Program, University of Pennsylvania, PA 19104
[6] Cell and Molecular Biology Graduate Program, University of Pennsylvania, PA 19104
[7] Chemical and Biomolecular Engineering Department, University of Pennsylvania, PA 19104
[8] Department of Chemistry, Texas A&M University, TX 77843
[9] Interdisciplinary Graduate Program in Genetics and Genomics, Texas A&M University, TX 77843
[10] Department of Bioengineering, University of Pennsylvania, PA 19104
[#] Equal contribution

**Supporting Information Includes:**

Title Page ------------------------ (Page S-1)
Extended Data Figure S1 ---- (Page S-2)
Extended Data Figure S2 ---- (Page S-3)
ED Figure S2 (continued) ---- (Page S-4)
Extended Data Figure S3 ---- (Page S-5, S-6)
Extended Data Figure S4 ---- (Page S-7)
Extended Data Figure S5 ---- (Page S-8)
Extended Data Table S1 (Page S-9)
Supplemental Figure S1 and text---- (Page S-10, S-11)

### Extended Data Figure S1

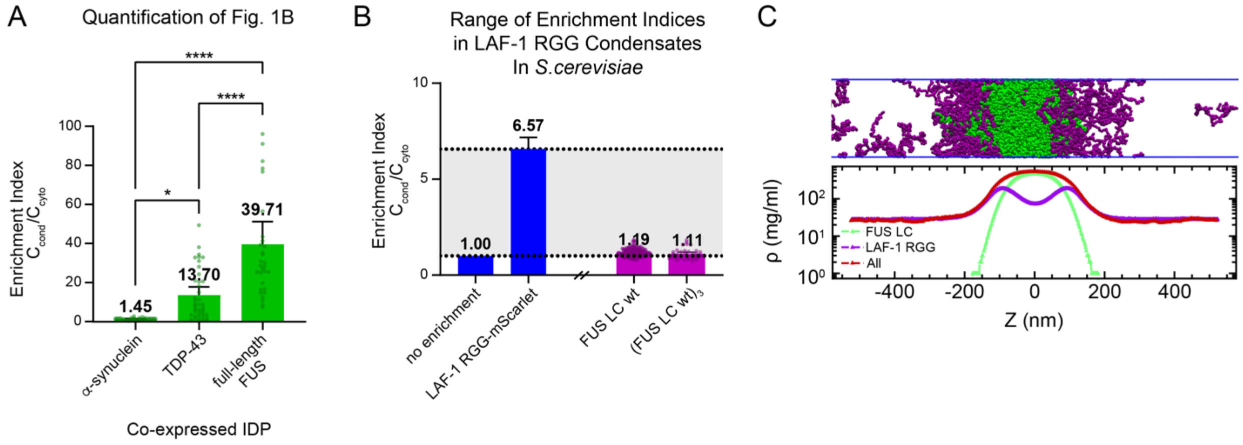


**Extended Data Fig. S1. Supplemental data for Figure 1.** A. Quantitation of IDP enrichment in LAF-1 RGG condensates from IDP co-expression screening in yeast; images from Fig 1B. EI values for 3 IDPs in (LAF-1 RGG)$_2$-mScarlet condensates. n ≥ 31 condensates per column. B. Plot showing possible range of enrichment indices in LAF-1 RGG condensates. Minimum value of 1. Maximum value based on LAF-1 RGG mScarlet (client) co-partitioning to (LAF-1 RGG)$_2$-GFP condensates. n ≥ 37 condensates per column. C. Schematic of CG slab at 300K consisting of FUS LC (green) and LAF-1 RGG (magenta). Distinct phases are observed with FUS LC forming a condensed phase and LAF-1 RGG sticking at its interface. Significance was calculated by one-way analysis of variance (ANOVA); ns P > 0.05, * P ≤ 0.05, ** P ≤ 0.01, *** P ≤ 0.001, and **** P ≤ 0.0001. Data relevant to Figure 1.

### Extended Data Figure S2

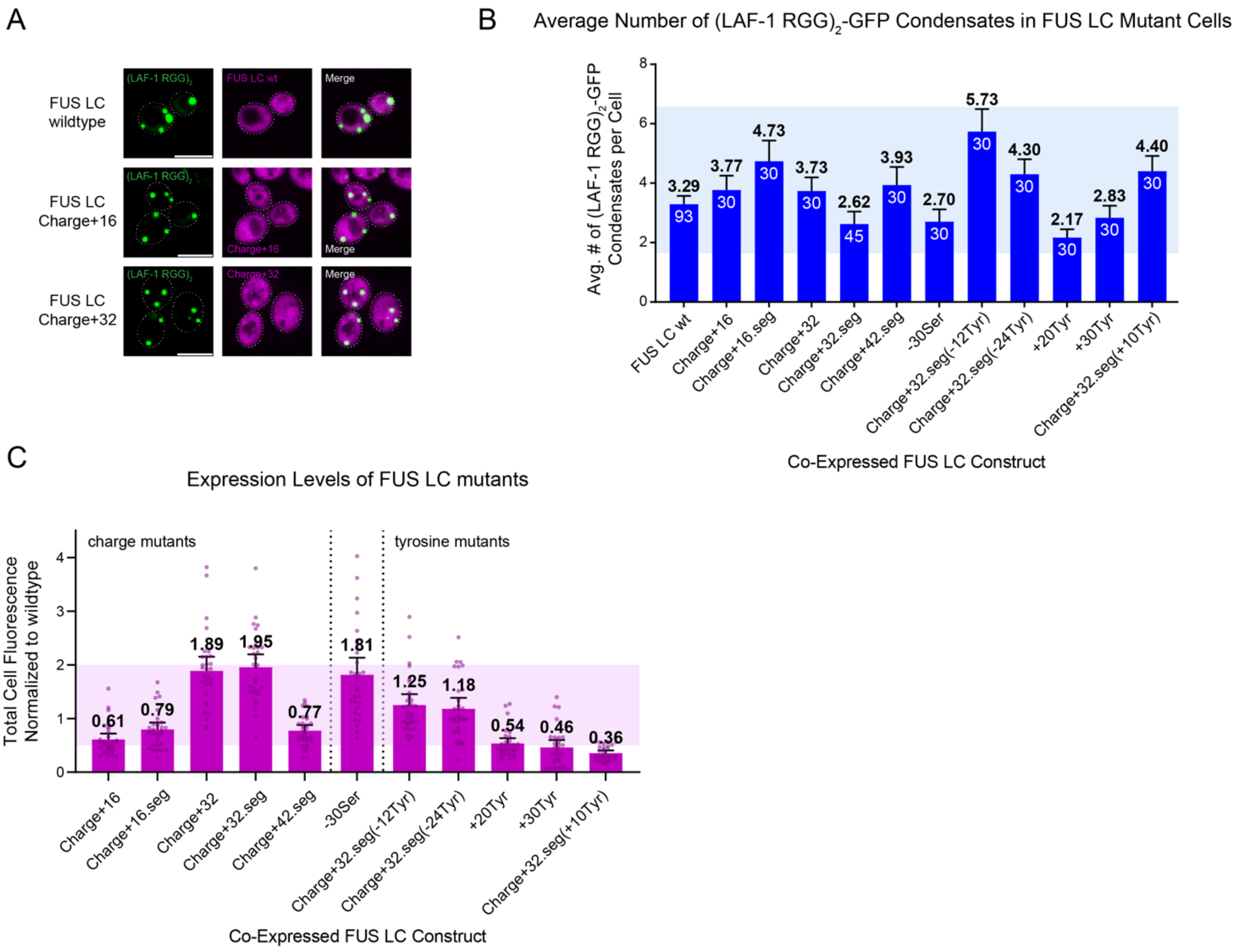


**Extended Data Fig. S2. Additional Images and controls for FUS LC Mutant Analysis.** A. Non-mixing behavior of FUS LC wildtype persists in mutants Charge+16 and Charge+32 when adding balanced charge. B. Average numbers of (LAF-1 RGG)$_2$-GFP condensates in co-expressing FUS LC mutant strains are within twofold of (LAF-1 RGG)$_2$ and FUS LC wildtype strain. Light blue box indicates range within twofold of wildtype. n ≥ 30 cells per column noted below column mean. C. Expression of FUS LC mutant constructs within co-expressed strains, normalized to FUS LC wildtype expression on day of imaging. Light magenta box indicates range within twofold of wildtype. n ≥ 30 cells per column. Error bars represent 95% confidence intervals. Significance was calculated by one-way analysis of variance (ANOVA); ns P > 0.05, * P ≤ 0.05, ** P ≤ 0.01, *** P ≤ 0.001, and **** P ≤ 0.0001. Scale bar: 5 μm. Data relevant to Figure 2.

### Extended Data Figure S2 (Continued)

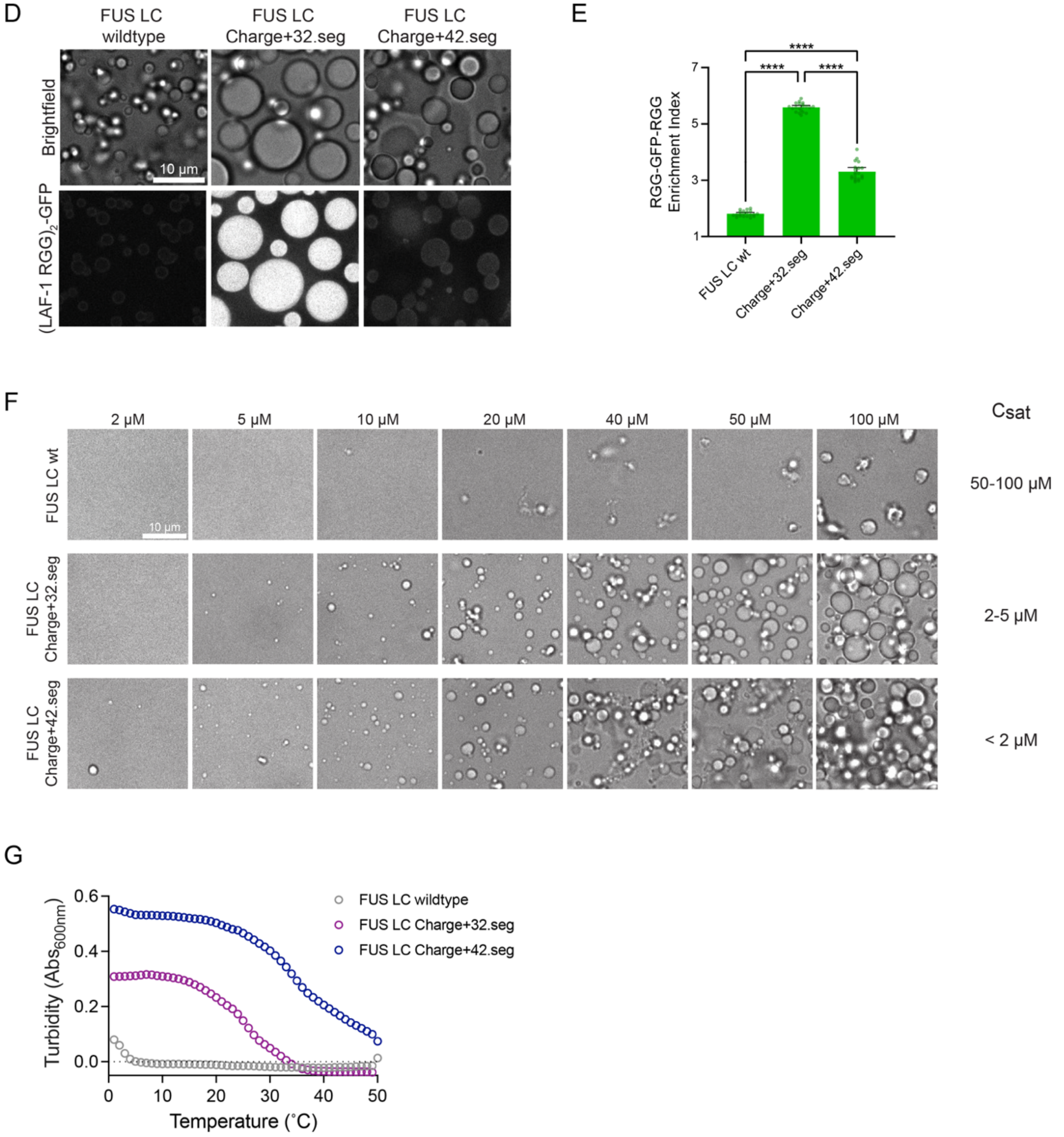


**Extended Data Fig. S2 (continued). LLPS specificity of purified FUS LC mutants and LAF-1 RGG proteins from biochemical reconstitution.** D. Partitioning of GFP-tagged (LAF-1 $RGG)_2$ to FUS LC wildtype and condensates in vitro**.** E. Quantitation of enrichment of (LAF-1 $RGG)_2$-GFP in condensed phase versus continuous phase from images in (D) 10 minutes after addition of (LAF-1 $RGG)_2$-GFP mutants to pre-formed FUS LC condensates (50 μM of each FUS LC construct). n = 20 condensates per column. F. Brightfield images of wells containing various concentrations of FUS LC mutants, showing their phase boundaries. Estimated $C_{sat}$ (right). G. Turbidity assays of FUS LC mutants (average of four independent trials) showing increased transition temperatures compared to wildtype. Error bars represent 95% confidence intervals. Scale bar: 10 μm. Data relevant to Figure 2.

### Extended Data Figure S3

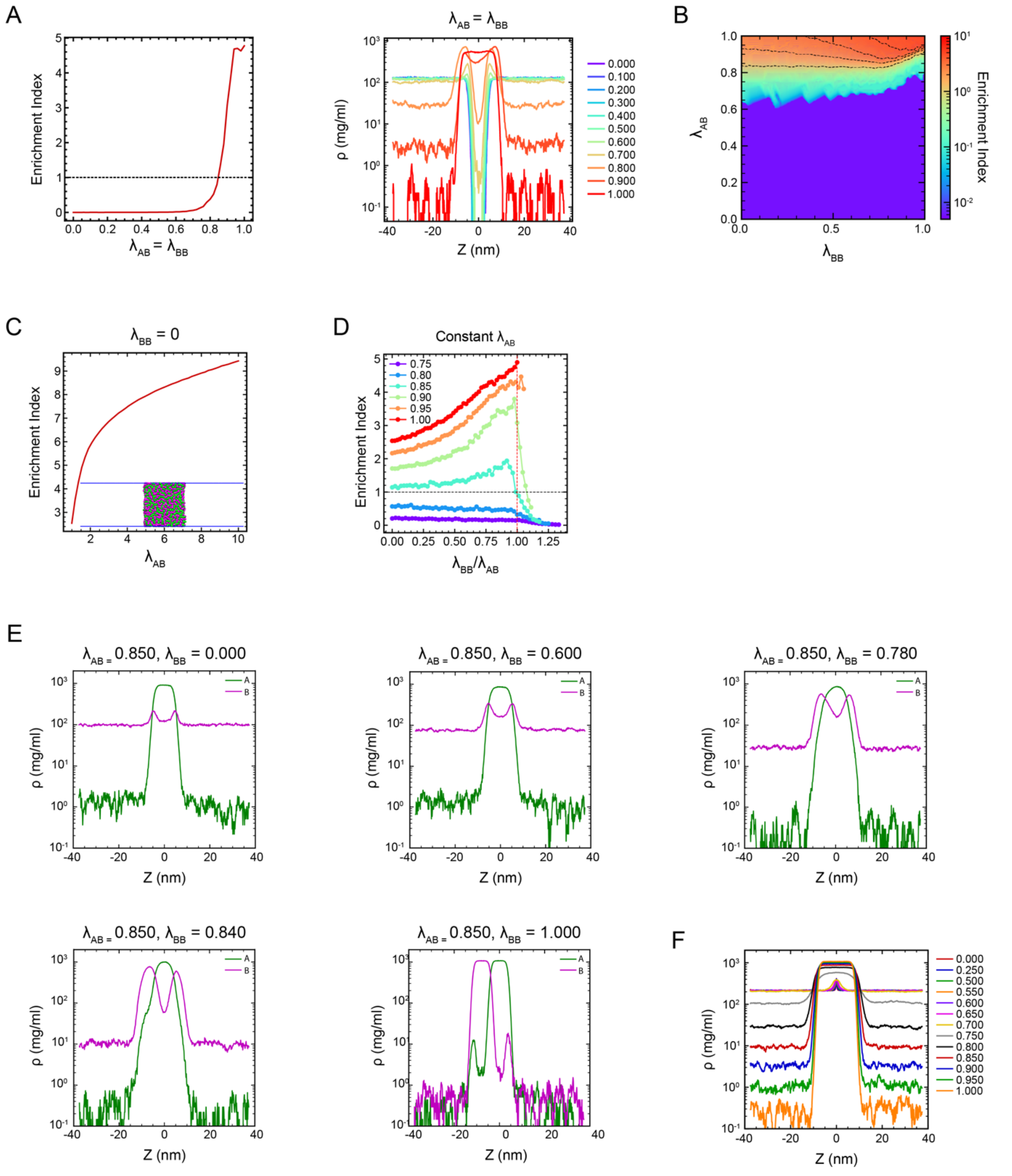


**Extended Data Fig. S3. Additional information from polymer model simulations.** A. Density profiles of polymer B and EI plot for $\lambda_{AB} = \lambda_{BB}$ case. B. Colormap of EI from CG scanning simulation with varying $\lambda_{AB}$ and $\lambda_{BB}$. C. Plot showing variation of EI as a function of $\lambda_{AB}$ at $\lambda_{BB}$ = 0 highlights the complex coacervate case. D. Plot showing variation of EI as a function of $\lambda_{BB}/\lambda_{AB}$ at different constant $\lambda_{AB}$ values. Maximum enrichment is observed when ratio of $\lambda_{BB}/\lambda_{AB}$ is close to 1. E. Density profiles of polymer A and B corresponding to the cartoons in Fig. 3E. F. Density profile from single component CG slab for varying interaction strength. For interaction strength for 0.75 or above, chains start to condense. Data relevant to Figure 3.

### Extended Data Figure S3 (Continued)

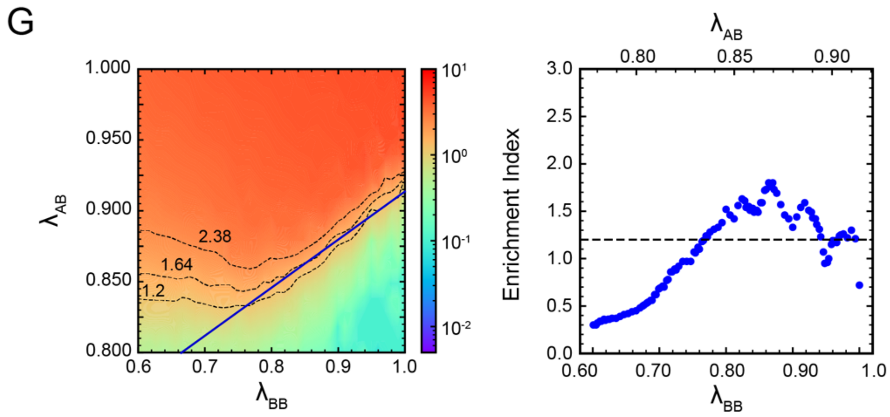


**Extended Data Fig. S3 (continued).** G. Simultaneous changes in homotypic and heterotypic interaction parameters can yield experimentally observed plateauing in EI. Plot shows variation of EI (right) along an arbitrarily defined change in $\lambda_{AB}$ and $\lambda_{BB}$ values (blue line shown in heat map (left)). Right plot shows plateauing of EI with increasing $\lambda_{BB}$ (bottom x-axis) and $\lambda_{AB}$ (upper x-axis). For +20Tyr and +30Tyr, increasing tyrosine will increase both homotypic and heterotypic interactions and scanning simulation shows one can obtain similar EI values as the FUS LC wildtype. For clarity, we only show one representative combination of $\lambda_{AB}$ and $\lambda_{BB}$, but there are several other possible combinations that can yield similar EI values as the FUS LC wildtype. Data relevant to Figure 3.

### Extended Data Figure S4

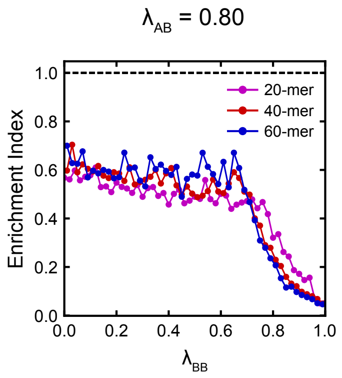


**Extended Data Fig. S4. Results from simulations varying polymer lengths.** Variation of EI with B's homotypic interaction at 0.80 heterotypic interaction. At low heterotypic interaction (≤ 0.80) longer chain-length do not enhance co-partitioning. Data relevant to Figure 3.

### Extended Data Figure S5

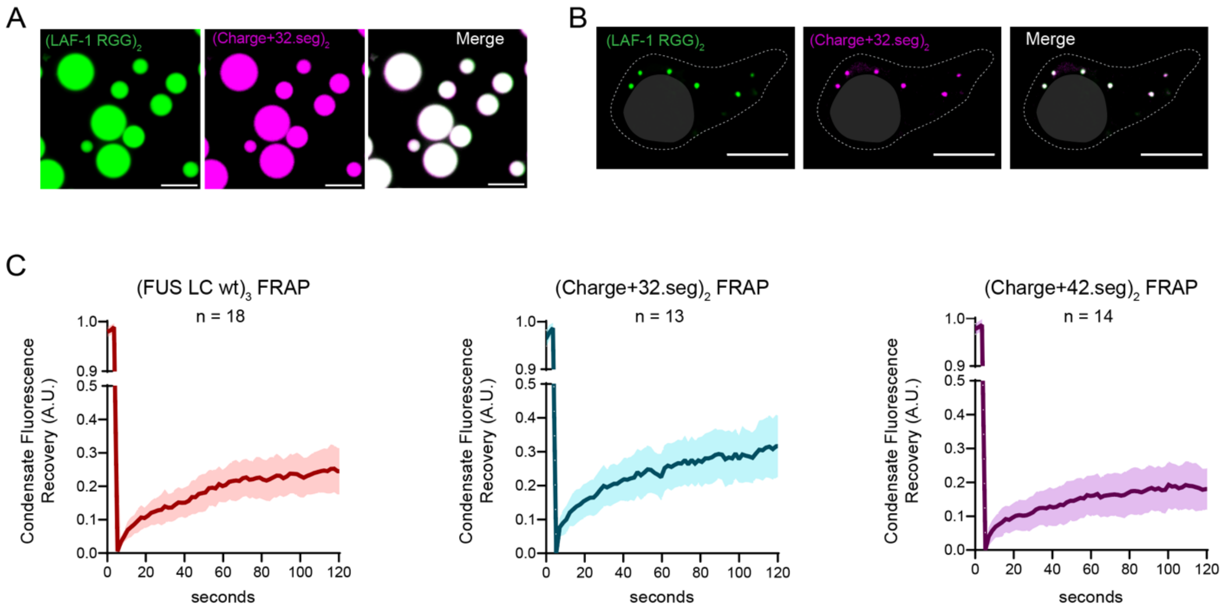


**Extended Data Fig. S5. Additional images relevant to LLPS specificity and formation of orthogonal condensates in vitro and in mammalian cells.** A. Biochemical reconstitution in vitro using purified proteins at concentrations above their $C_{sat}$, to generate droplets. (LAF-1 RGG)$_2$-GFP strongly co-partitions with FUS LC Charge+32.seg condensates; (FUS LC)$_2$-BFP tracer; "merge" in white shows overlay. B. Loss of LLPS specificity for FUS LC mutant charge+32.seg in HEK293T cells co-transfected with (LAF-1 RGG)$_2$-GFP; (FUS LC Charge+32.seg)$_2$-mCherry strongly co-partitions with (LAF-1 RGG)$_2$-GFP condensates. Scale bar: 10 $\mu$m. C. FRAP plots from condensates of FUS LC constructs when expressed as multimers. Left: Triple version of FUS LC wildtype. Middle: Tandem version of Charge+32.seg. Right: Tandem version of Charge+42.seg. Plots show 95% confidence intervals. Samples sizes are shown on each plot. Data relevant to Figure 5.

### Extended Data Table S1

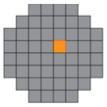 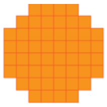

| FUS LC construct | Max pixel enrichment index | | Average pixel enrichment index | |
|---|---|---|---|---|
| | mean | 95% confidence interval | mean | 95% confidence interval |
| FUS LC wildtype | 1.19 | ±0.033 | 1.11 | ±0.022 |
| Charge+16 | 1.16 | ±0.042 | 1.08 | ±0.025 |
| Charge+16.seg | 1.25 | ±0.036 | 1.11 | ±0.023 |
| Charge+32 | 1.25 | ±0.037 | 1.11 | ±0.025 |
| Charge+32.seg | 1.64 | ±0.082 | 1.30 | ±0.035 |
| -30Ser | 1.19 | ±0.066 | 1.12 | ±0.046 |
| Charge+42.seg | 2.38 | ±0.168 | 1.56 | ±0.063 |
| Charge+32.seg(-12Tyr) | 1.21 | ±0.034 | 1.12 | ±0.024 |
| Charge+32.seg(-24Tyr) | 1.08 | ±0.026 | 1.05 | ±0.020 |
| Charge+32.seg(+10Tyr) | 1.71 | ±0.103 | 1.42 | ±0.080 |

**Extended Data Table S1. Enrichment indices of FUS LC clients using mean client values.** The second and third columns show the FUS LC client EIs as calculated throughout the manuscript using the max 488 nm pixel in each channel as the peak value in EI calculation (along with 95% confidence intervals). The fourth and fifth columns show values if EIs were instead calculated by averaging all 561 nm pixel values within the thresholded LAF-1 RGG condensate, frequently averaging in artificially low values due to variability in signal (also with 95% confidence intervals). Due to low expression, mutants +20Tyr and +30Tyr were analyzed using a different technique and could not be quantified with this alternative procedure (see Methods).

### Supplemental Text 1. Simulations with angle constraints

We conducted additional scanning simulations to examine the role of configurational entropy in our model. In our minimalistic model, we considered polymer A as the scaffold and polymer B as the client, mimicking the experimental system. Both polymers were initially flexible without any angle constraints. To investigate the influence of configurational entropy, we introduced rigidity to polymer B while keeping polymer A flexible. The rationale behind this was to lower the configurational entropy of polymer B and observe its impact on the system. We hypothesized that by making polymer B rigid, its configurational entropy would decrease, leading to an early enrichment of polymer B within polymer A. We didn't have a clear hypothesis about how it will change the turnover point and enrichment beyond this point To make the protein chains rigid, we introduced angle constraints between three consecutive monomers in polymer B. Specifically, we set an equilibrium angle of 180 degrees and varied the angle constant ($k_\theta$) to different values (0.5, 1, 2, 3, 4, 5, 10 kcal/mol*rad^2). These values were selected based on the single chain simulations to span different chain rigidity values (Supplementary Fig. 1 A,B).

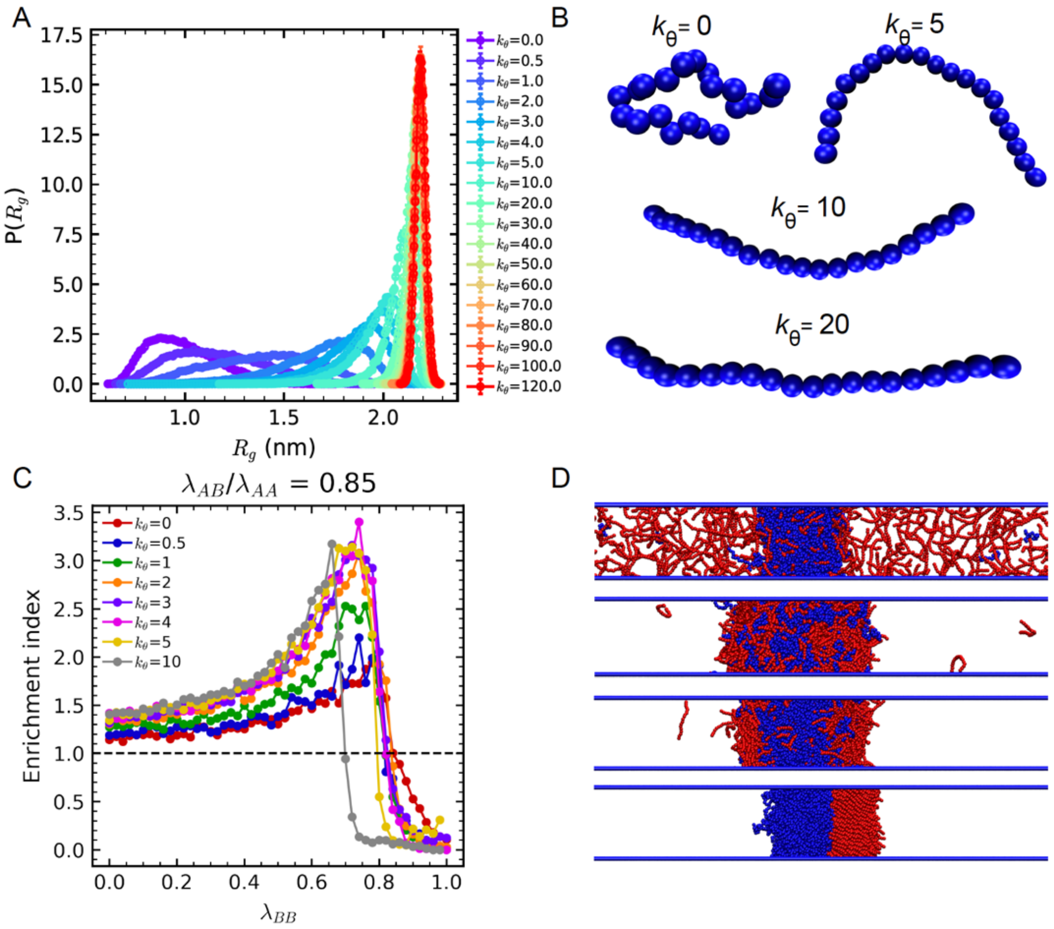


**Supplementary Fig. 1: Rigid polymer simulation.** A. Probability distribution of radius of gyration $R_g$ from single chain simulations for different angle constant ($k_\theta$). B. Cartoon representation of single chain at different rigidity. C. Variation in EI as a function of $\lambda_{BB}$ at a constant $\lambda_{AB}$ (0.85) for different angle constants. D. Cartoon representations of two component system with flexible polymer A (blue) and rigid polymer B (red). Cartoons from top to bottom represents the scenarios with increasing $\lambda_{BB}$ at a constant $\lambda_{AB}$.

Interestingly, we observed a similar non-monotonic trend for both rigid and flexible chains at a constant $\lambda_{AB}$ value of 0.85 (Supplementary Fig 1C). The lower configurational entropy of the rigid polymers ($k_\theta > 0$) make these partition inside the condensed phase at lower $\lambda_{BB}$ values. Furthermore, as the angle constant increased, we noticed that the crossover point at which the enrichment started to decrease shifted to lower $\lambda_{BB}$ values, presumably due to the higher tendency of rigid B polymers to form homotypic condensed phase (Supplementary Fig 1D).

Overall, our simulation results demonstrate that the model effectively captures the influence of configurational entropy, with the equilibrium behavior ultimately dictated by the free energy minimum, which is the central goal of this study.
